# Heterotypic electrostatic interactions control complex phase separation of tau and prion into multiphasic condensates and co-aggregates

**DOI:** 10.1101/2022.09.17.508370

**Authors:** Sandeep K. Rai, Roopali Khanna, Anamika Avni, Samrat Mukhopadhyay

## Abstract

Biomolecular condensates formed via phase separation of proteins and nucleic acids are thought to perform a wide range of critical cellular functions by maintaining spatiotemporal regulation and organizing intracellular biochemistry. However, aberrant phase transitions are implicated in a multitude of human diseases. Here, we demonstrate that two neuronal proteins namely, tau and prion undergo complex coacervation driven by domain-specific electrostatic interactions to yield highly dynamic, mesoscopic liquid-like droplets. The acidic N-terminal segment of tau interacts electrostatically with the polybasic N-terminal intrinsically disordered segment of the prion protein (PrP). We employed a unique combination of time-resolved tools that encompass several orders of magnitude of timescales ranging from nanoseconds to seconds. These studies unveil an intriguing orchestra of molecular events associated with the formation of heterotypic condensates comprising ephemeral, domain-specific, short-range electrostatic nanoclusters. Our results reveal that these heterotypic condensates can be tuned by RNA in a stoichiometry-dependent manner resulting in reversible, multiphasic, immiscible, ternary condensates of different morphologies ranging from core-shell to nested droplets. This ternary system exhibits a typical three-regime phase behavior reminiscent of other membraneless organelles including nucleolar condensates. We also show that upon aging, tau-PrP droplets gradually convert into solid-like co-assemblies by sequestration of persistent intermolecular interactions. Our vibrational Raman spectroscopic data in conjunction with atomic force microscopy and multi-color fluorescence imaging results reveal the presence of amorphous and amyloid-like co-aggregates upon maturation. Our findings provide mechanistic underpinnings of overlapping neuropathology involving tau and PrP and highlight a broader role of complex phase transitions in physiology and disease.

## Introduction

Biomolecular condensates formed via liquid-liquid phase separation of proteins and nucleic acids serve as on-demand membraneless organelles that are involved in the spatiotemporally-controlled organization, compartmentalization, and regulation inside living cells. Owing to their ability to form and dissipate in response to cellular cues, regulatability, permeability, and the ability to selectively concentrate the biomolecules, these noncanonical liquid-like organelles are emerging as central players at all levels of essential cellular activity ranging from the expression and regulation of genes to the modulation of intricate signaling pathways (1-11). An emerging body of work has revealed that intrinsically disordered proteins/regions (IDPs/IDRs) often comprising polypeptide repeat units, low sequence complexity, and prion-like domains are ideal candidates for biological phase separation. Such sequence features offer a fuzzy network of weak, multivalent, non-covalent, and transient contacts that govern the relay of making and breaking of weak interactions and promote phase separation into liquid-like compartments (12-19). The protein phase behavior and the internal material property can be tuned by other proteins and nucleic acids such as RNA. Heterotypic interactions between the molecular entities involving a multitude of proteins and nucleic acids can often lead to the formation of multi-component, multiphasic, and mutually immiscible biomolecular condensates such as nucleolar condensates (20-25). In vitro, many of the essential biophysical features of liquid-like intracellular membraneless organelles can be recapitulated by using purified proteins with or without nucleic acids that can spontaneously demix from a mixed homogeneous phase into two co-existing phases namely, the dense phase and the light phase (11, 26, 27). The current proposal of biomolecular condensation involves a density transition coupled to percolation that results in the condensed phase comprising a viscoelastic network fluid (4). Such physical microgels can undergo maturation via time-dependent changes in the material property into solid-like aggregates. These protein aggregates formed via aberrant phase transitions are thought to be involved in a range of deadly neurodegenerative diseases such as Alzheimer’s Disease, Amyotrophic Lateral Sclerosis (ALS), Frontotemporal Lobar Degeneration (FTLD), and so forth (28-32). Therefore, the understanding of the precise molecular determinants of biological phase separation is of great importance in both physiology and pathology. In the present work, we describe an intriguing interplay of two neuronal proteins namely, tau and prion proteins that undergo complex coacervation into heterotypic condensates. These condensates transform into multiphasic condensates in the presence of RNA and exhibit a time-dependent maturation into solid-like aggregates.

Tau is a microtubule-associated neuronal IDP. The human brain expresses six different spliced variants (33, 34). The longest isoform, full-length tau, harbors two N-terminal inserts and a proline-rich domain followed by four repeat regions, one pseudo repeat region, and the C-terminal end (Fig. 1*A*). Under normal conditions, tau interacts with microtubular proteins mediated by the proline-rich domain and the repeat region. These regions are prone to modifications in the form of posttranslational hyperphosphorylation under diseased conditions (35, 36). Additionally, the tau chain contains a heterogeneous cluster of charged residues, oppositely charged domains, and polar residues with a high proline and glycine content throughout the sequence, making it a highly dynamic and amphipathic polypeptide. Recent reports have shown that under physiological and near-physiological conditions, tau can undergo a liquid-to-solid transition via phase separation driven by homotypic as well as heterotypic interactions in the absence or presence of a crowding agent (37-40). Such condensates have been proposed to act as crucibles for aberrant phase transitions involved in disease progression. Tau pathology extends to many other neuronal and RNA-binding proteins (RBPs). The interactions of tau and its hyperphosphorylated variants with several RBPs, including Musashi and T-cell intracellular antigen 1, have been proposed to result in heterotypic inclusions that might be responsible for the exacerbation of overlapping neuropathological features and diseases (41). Moreover, the accumulation of neurofibrillary tangles (NFTs) of tau with another IDP, α-synuclein (α-Syn), in Parkinson’s disease hints toward the synergistic interactions between tau and α-Syn (42). Tau protein deposits have also been found in the brains of patients affected by the prion protein cerebral amyloid angiopathy (PrP-CAA). These patients displayed a nonsense mutation (Q160Stop) in the *PRNP* gene that generates a highly unstructured variant of this polypeptide chain due to translation termination. Additionally, Gerstmann-Sträussler-Scheinker (GSS) syndrome is also attributed partly to tau deposits, where a missense mutation in the *PRNP* gene leads to an amino acid switch at residue 198 (F198S) of PrP (43-45). The cellular form of PrP (PrP^C^) is a GPI-anchored protein consisting of an N-terminal signal peptide (residues 1-23), a highly positive disordered N-terminal tail containing oligopeptide repeats (residues 23-120), a globular C-terminal domain (residues 121-230), and a GPI-anchor signal (residues 231-253) (46-48) (Fig. 1*B*). It also contains a putative RNA binding site and is often classified as an RBP. As a step toward elucidating the overlapping neuropathological features of the two proteins namely, tau and PrP, we set out to investigate the role of PrP in regulating the phase behavior of tau and discovered an intriguing interplay of molecular drivers in modulating complex phase transitions.

**Fig 1.**
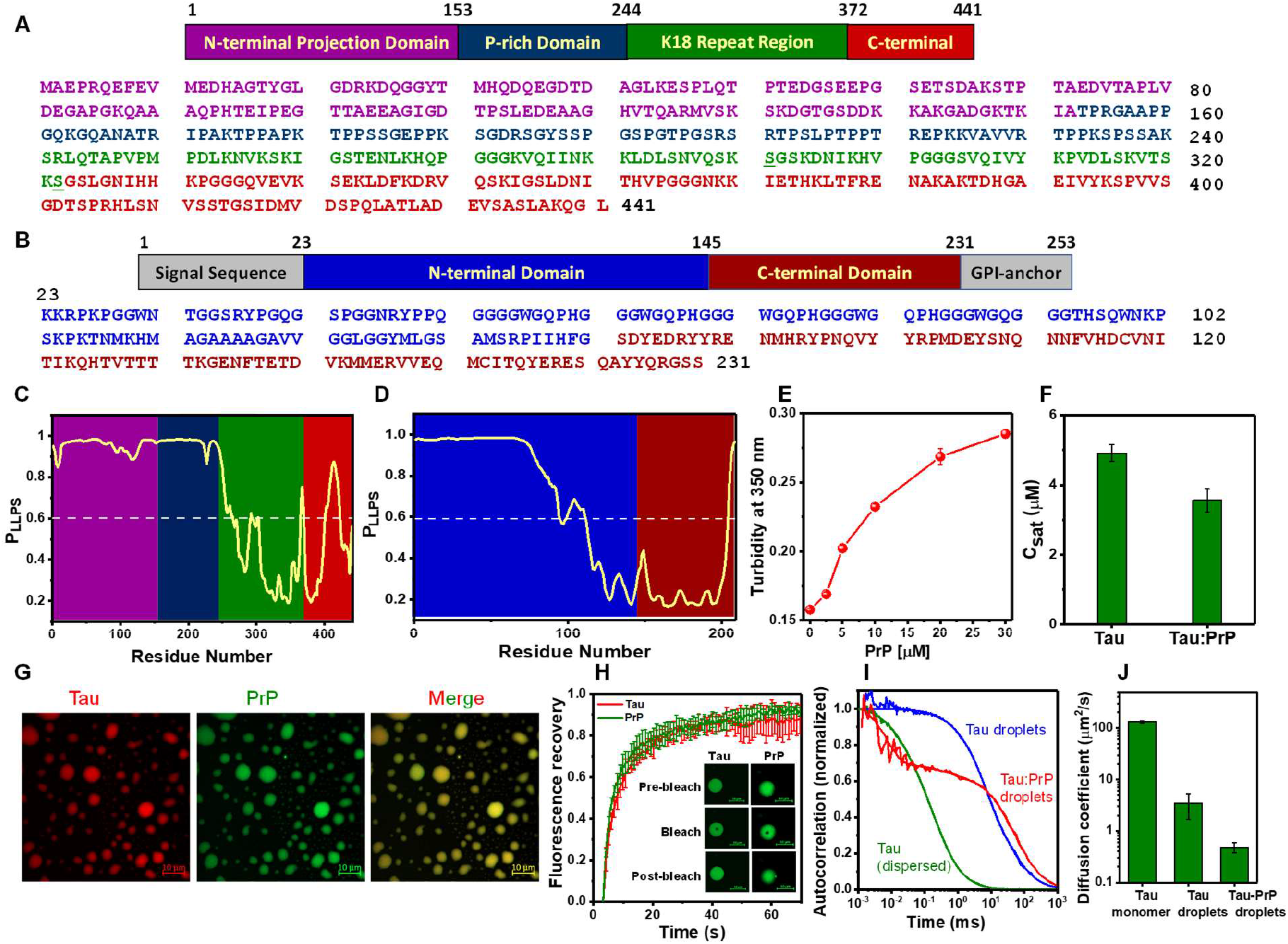
PrP induces phase separation of tau. **(A)** Domain architecture and amino acid sequence of full-length tau. **(B)** Different segments and amino acid sequence of full-length PrP (23-231). **(C)** The propensity of phase separation predicted for tau and **(D)** PrP using FuzDrop. **(E)** The turbidity plot of tau (10 μM) in the presence of an increasing concentration of PrP. **(F)** Saturation concentration (C_sat_) estimated by ultracentrifugation. **(G)** A high-resolution two-color Airyscan image of colocalized tau (tau T17C-Alexa594-labeled, red) and PrP (PrP W99C-Alexa488-labeled, green) in tau:PrP coacervates (scale bar 10 μm). Tau and PrP concentrations were 10 and 20 μM, respectively. The imaging was performed at least thrice with similar observations. **(H)** FRAP kinetics of tau:PrP droplets. Alexa488-labeled proteins were used for the FRAP for both tau and PrP. The data represents mean ± s.d.; n= 5. **(I)** The normalized autocorrelation plot obtained from FCS measurements performed for 5 different droplets. The unnormalized autocorrelation plots are shown in Fig. S1. **(J)** Diffusion coefficients of tau in monomeric dispersed form, tau-only droplets, and tau:PrP droplets obtained from FCS. measurements (n = 5). For all FCS measurements, 5 nM Alexa488-labeled tau T17C was used. Tau and PrP concentrations were 10 and 20 μM, respectively.

## Results

### PrP promotes spontaneous phase separation of tau through heterotypic interactions

Sequence composition governs the phase behavior of a protein by tuning its multivalent interactions. Clustered negatively and positively charged residues, weakly hydrophobic segments, and cation-π modulators such as arginine and aromatic amino acids are important for regulating these interactions. We first analyzed the full-length Tau protein. It has a negatively charged N-terminal and positively charged Proline-rich domain, in addition to a hydrophobic repeat domain and a mildly negatively charged C-terminal end, making it a highly unstructured amphipathic polymer and hence an ideal candidate for phase separation (Fig. 1*A*). This was further supported by sequence-based predictors such as FuzDrop (49) and catGRANULE (50), which predicted a high phase separation propensity for the tau protein (Fig. 1*C* and Fig. S1*A*). We began with turbidity measurements and confocal microscopy to complement our bioinformatic analyses with experimental proof and establish the in vitro phase separation of tau at a near-physiological condition. Under our condition at pH 6.8 (14 mM HEPES), tau remains in a monomeric dispersed form even at very high concentrations in the presence of as low as 50 mM NaCl. Upon lowering the salt concentration (∼ 10 mM NaCl), the turbidity of the protein solution rose even at protein concentrations as low as ∼ 6 μM, which is close to the physiological concentration of tau (2 μM). To further verify the increase in turbidity with observable evidence, we used an AlexaFluor488-maleimide labeled single-cysteine variant of the protein (Tau T17C) to perform the confocal fluorescence imaging. Tau protein doped with a labeled variant (> 1%) was used for imaging, which reveals the formation of tau protein droplets under this condition (Fig. S1*B*). The observed droplets had a typical liquid-like behavior and, in addition to having a rapid fluorescence recovery after photobleaching (FRAP), also underwent characteristic fusions and surface-wetting (Fig. S1*D* and S1 video). These findings demonstrate the coacervation of tau via homotypic interactions and are in accordance with previous findings on the phase separation of tau (37, 38). Previous results have indicated the ability of RNA to act as an inducer for protein phase separation by engaging in heterotypic interactions with polypeptide chains. Under our conditions, a low concentration of RNA (∼ 5 ng/μL) was able to enhance the phase-separation ability of tau as evinced by increased turbidity and microscopy imaging (Fig. S1*B*). These tau:RNA complex condensates behave similarly to tau-only droplets exhibiting liquid-like behavior and rapid FRAP (Fig. S1*D)*.

We next investigated the effect of PrP on the phase behavior of tau. Of note here is that, though PrP showed a high phase separation tendency based on predictor (Fig. 1*D*), it does not undergo spontaneous phase separation under our conditions (Fig. S1*C*). To this end, we measured the turbidity of tau in the presence of an increasing concentration of PrP. Our data showed an increase in the turbidity upon the addition of PrP suggesting the increase in the phase separation propensity of tau (Fig. 1*E*). To quantify this observation, we performed ultracentrifugation to estimate and compare the saturation concentration (C_sat_) of tau-only and tau:PrP droplets. These results indicated the lowering of C_sat_ values in the presence of PrP (Fig. 1*F*). We next performed two-color confocal imaging to visualize the complex coacervation of tau and PrP. We labeled single-Cys variants of tau (T17C) and PrP (W99C) using AlexaFluor594 maleimide and Alexa Fluor488 maleimide, respectively. We carried out phase separation assays with the tau and PrP in the presence of ∼ 1% labeled proteins and imaged them under a confocal microscope. Two-color imaging revealed the colocalization of tau and PrP within these liquid condensates (Fig. 1*G*). Complex coacervation of tau and PrP yielded a much larger number of droplets that were smaller and more spherical compared to tau-only droplets (Fig. S1*F* and S2 video). Next, in order to delineate the material properties of these droplets, we performed FRAP experiments that revealed rapid near-complete recovery for both proteins indicating their high mobility inside the droplets (Fig. 1*H*). Further to probe the properties of the dense phase on a microsecond to sub-millisecond timescale, we monitored the diffusion time of Alexa Fluor488 maleimide labeled tau protein inside tau only and tau:PrP droplets by fluorescence correlation spectroscopy (FCS). The Diffusion time extracted from our correlation measurements suggested the presence of stronger intermolecular interactions in the case of tau:PrP droplets, i.e. stronger networking, in comparison to tau-only droplets (Fig. 1*I* and Fig. S1*G-I*). Taken together, our observations suggest that PrP potentiates the phase separation of tau and is recruited within tau droplets resulting in a strongly inter-connected chain network. Next, to further unmask the nature of intermolecular interactions driving tau:PrP complex coacervation, we investigated the sequence features of both proteins. Given the clustering of opposite charges in both proteins, we hypothesized that they might interact electrostatically in a manner similar to what has been reported before (51, 52). To investigate this, we set out to delineate the effect of salt on tau:PrP complex coacervation.

### Electrostatic interactions are a principal driver of tau-PrP heterotypic coacervation

Many oppositely charged polypeptides, proteins, and synthetic polyelectrolytes undergo stoichiometry-dependent charge neutralization that drives their phase separation upon mixing highlighting the central role of electrostatic interactions in this process. Tau and PrP are similarly charged biopolymers having an isoelectric point (pI) of 8.24 and 9.44, respectively. Based on their overall charge, one would expect repulsion between these two chains. However, upon closer inspection of the amino acid sequence and charge distribution throughout both protein chains (53), we found that tau and PrP chains are oppositely charged at the N-terminal ends which can drive their heterotypic assembly (Fig. S2*A* and S2*B*). To test our hypothesis, we performed salt-dependent turbidity assays and confocal imaging of the tau:PrP condensates (Fig. 2*A* and Fig. S2*C*). With an increasing salt concentration, the turbidity of the tau:PrP mixture dropped sharply with a dissolution of droplets at ∼ 50 mM NaCl highlighting the role of intermolecular electrostatic interactions driving the complex coacervation. With increasing PrP concentration, these heterotypic assemblies were able to sustain higher salt concertation (Fig. 2*B* and Fig. S2*C*, S2*D*). At a higher total protein concentration (tau 30 μM and PrP 60 μM), we observed phase separation even at a physiological salt concentration (150 mM) (Fig. S2*E*). Post-translational modifications (PTMs) of a protein often play a crucial role in modulating its phase separation ability and interactions with other proteins or biomolecules primarily by changing its charge distribution (54, 55). We next probed whether the frequently occurring phosphorylation of tau, which adds negative charges onto the tau chain, would increase the phase separation propensity of tau:PrP. To test this, we created a triple phosphomimetic variant of tau (tau 3P) by selectively mutating three residues to glutamate (S202E, S205E, T208E). This triple phosphomimetic variant of tau (tau 3P) exhibited a higher propensity for phase separation with PrP as observed by our turbidity assays (Fig. 2*C*). Further, even a much lower concentration of PrP (∼ 2 μM) promoted the phase separation of tau 3P compared to wild-type tau. Two-color imaging also corroborated our turbidity measurements (Fig. 2*D*). Imaging and FRAP experiments validated the liquid-like nature of tau 3P-PrP droplets (Fig. S2*F*). These tau 3P-PrP droplets were larger and more mobile compared to tau-PrP droplets as observed by the more frequent fusion events in the case of tau 3P-PrP (Fig. S2*G*). These results together suggested that electrostatic interactions modulate the complex coacervation of tau and PrP. We next set out to delineate the roles of specific protein domains in driving the complex coacervation of tau and PrP.

**Fig 2.**
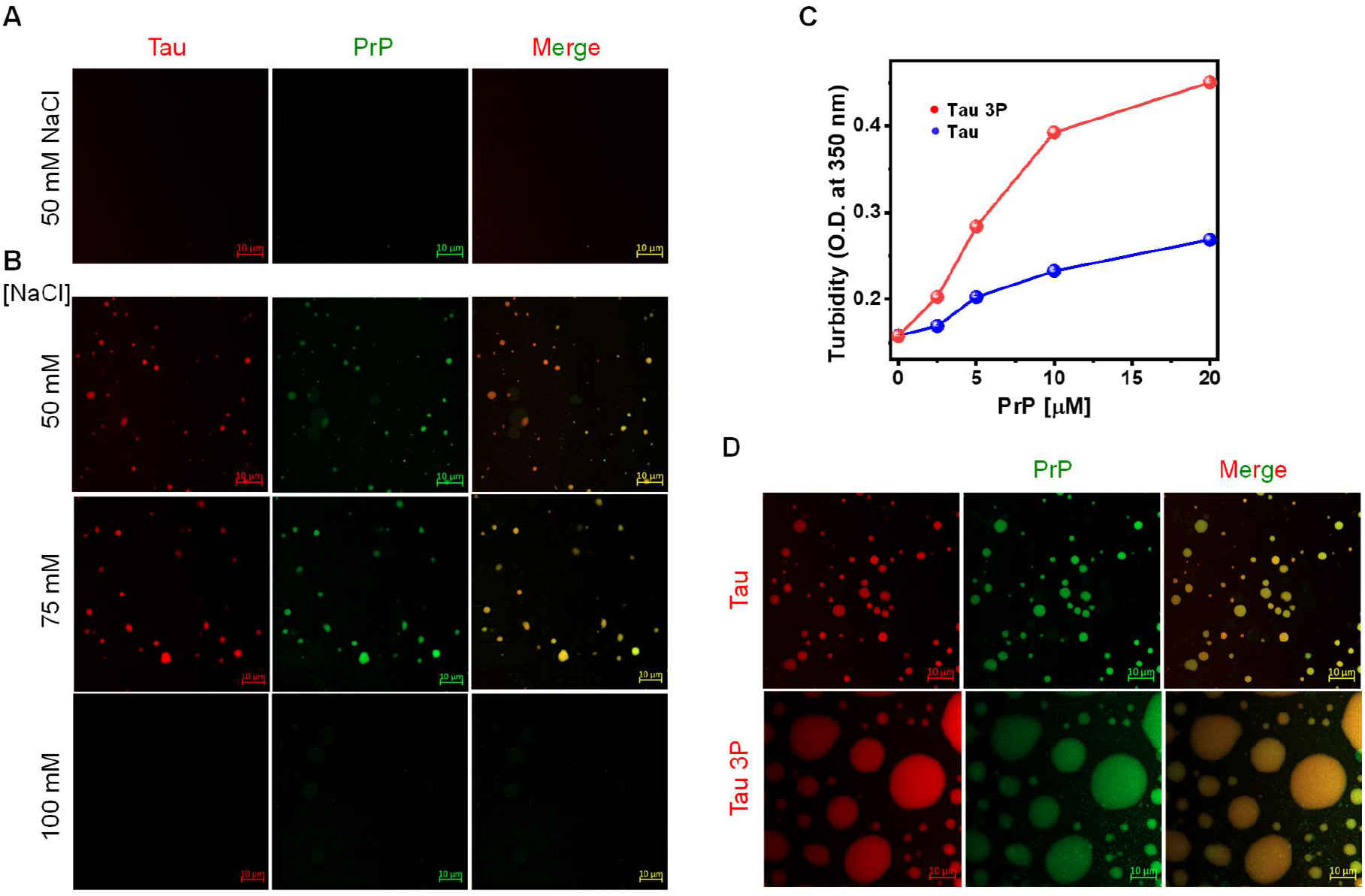
Electrostatic interactions govern tau-PrP phase separation. **(A)** Salt-induced dissolution of tau:PrP droplets in the presence of NaCl (scale bar, 10 μm). **(B)** Tau (10 μM, red) and PrP (50 μM, green) complex coacervates in the presence of an increasing concentration of salt (scale bar 10 μm). **(C)** The turbidity plot of tau and triple phosphomimetic mutant (tau 3P) against an increasing concentration of PrP. The data represent mean ± s.d.; n= 3. **(D)** Two-color high-resolution Airyscan images of tau 3P (10 μM, red) and PrP (10 μM, green) droplets (lower panel) compared with tau (upper panel) (scale bar, 10 μm).

### Domain-specific interactions drive the co-condensation of tau and PrP

In order to obtain domain-specific insights into the electrostatic interactions between tau and PrP, we created several truncation variants of both proteins (Fig. 3*A* and 3*B*). First, we aimed at characterizing the role of the negatively charged N-terminal fragment and positively charged P-rich region of tau and created two naturally occurring truncation variants namely, Nh2-tau (aa 26-230, pI = 5.32) and tau 0N4R (aa 151-391, pI = 10.23). These variants of tau are found in NFTs of tau deposits in the Alzheimer’s disease brain and are thought to play a vital role in tau pathogenesis (56, 57). Based on the high net negative charge, we posited that Nh2-tau (net charge ∼ - 5) could undergo phase separation with PrP (predicted net charge ∼ + 10) by a charge-neutralization mechanism that is reminiscent of RNA-driven reentrant phase transitions. Indeed, our turbidity measurements showed a reverse bell-shaped profile for Nh2-tau:PrP at a fixed concentration of PrP (Fig. 3*C*). The turbidity value peaked at a 2:1 stochiometric ratio of Nh2-tau:PrP where the charge neutralization was achieved. Our two-color confocal microscopy also corroborated this observation (Fig. 3*D*). These Nh2-tau:PrP droplets showed liquid-like characteristics, grew with time, and after 1 hr these droplets appear to completely coalesce (Fig. S3*A*). Further, as expected, the other positively charged fragment of tau, tau 0N4R, did not phase separate, either alone or in the presence of PrP (Fig. 3*E*). These results indicated that the negatively charged N-terminal domain of tau is important for heterotypic condensation of tau and PrP.

**Fig 3.**
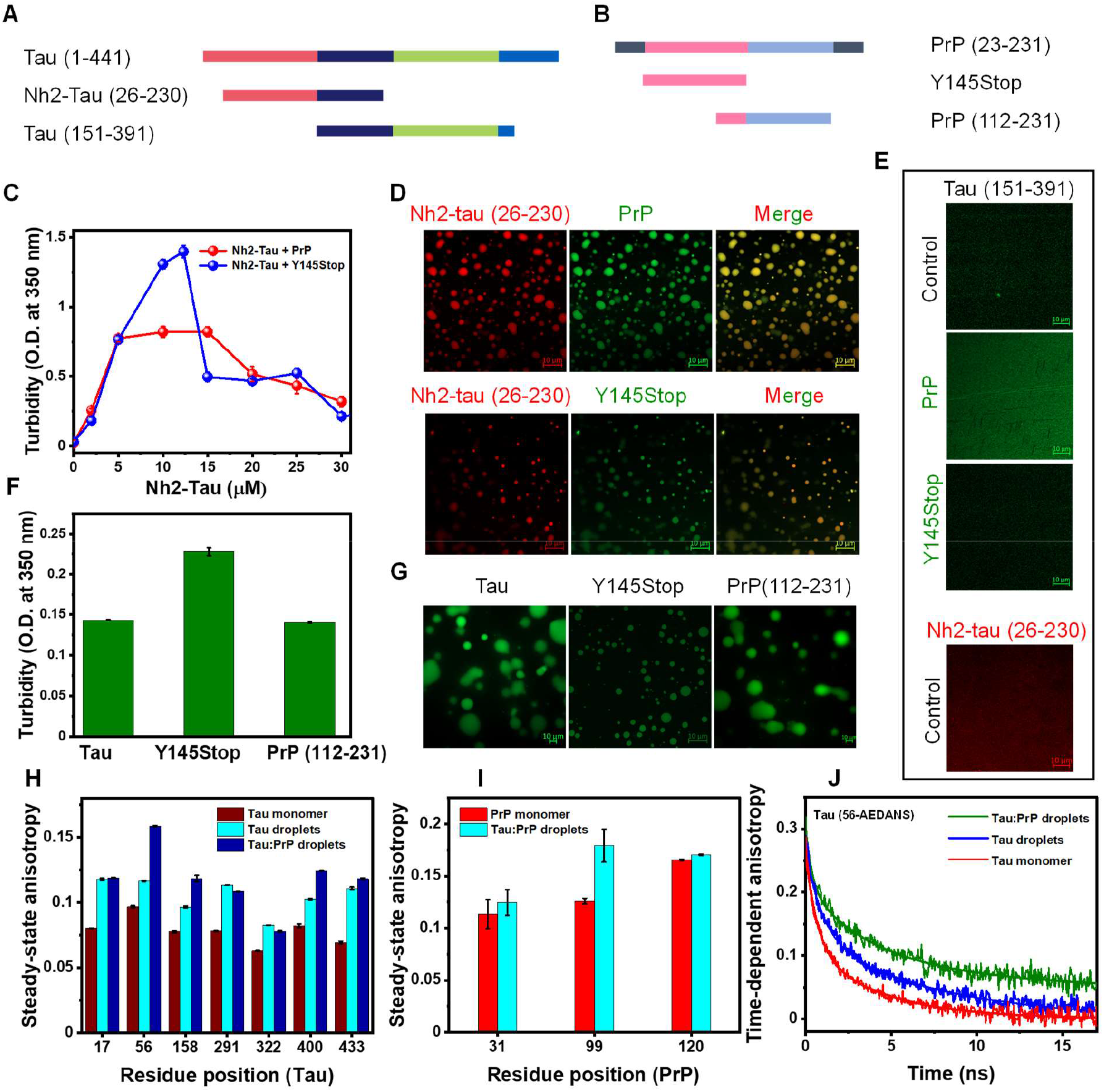
Heterotypic interactions between the N-terminal domains of tau and PrP drive complex coacervation. **(A-B)** The framework of tau and PrP truncations used. **(C)** The turbidity plot of Nh2-tau:PrP and Nh2-tau:Y145Stop. The data represent mean ± s.d.; n = 3. **(D)** Two-color confocal images of Nh2-tau:PrP (upper panel) and Nh2-tau:Y145Stop (lower panel) droplets (scale bar 10 μm). **(E)** Images of the mixed homogeneous phase of 0N4R tau (upper panel, green), 0N4R with PrP (second panel, green), 0N4R with Y145Stop (third panel, green), and Nh2-tau (bottom panel, red) (scale bar, 10 μm). **(F)** Effect of PrP truncations (20 μM) on tau (10 μM) turbidity. The data represent mean ± s.d.; n = 3. **(G)** Confocal images of tau in the presence of PrP truncations (scale bar 10 μm). **(H)** Steady-state fluorescence anisotropy measurements of F-5-M labeled single Cys mutants of tau, spanning the complete sequence, in the monomeric (wine), tau-only droplet (cyan), and tau:PrP coacervates (blue) forms. The data represent mean ± s.d.; n = 3. **(I)** Steady-state fluorescence anisotropy measurements of F-5-M labeled single Cys mutants of PrP in the monomeric (red) and tau:PrP coacervate (cyan) forms. The data represent mean ± s.d.; n = 3. **(J)** Time-resolved fluorescence anisotropy decays of IAEDANS labeled at S56C of tau in monomer (red), tau-only droplets (blue), and tau:PrP droplets (olive). The solid lines are fits obtained using decay analysis for monomers and droplets, respectively. Similar anisotropy decay profiles were obtained for the N-terminal segment of PrP (Fig. S3*H*). See SI for details of measurements and analysis and Table S2 for recovered parameters.

In order to investigate the role of different regions of PrP in tau:PrP co-condensation, we used a naturally occurring, pathological C-terminally truncated variant of PrP namely, Y145Stop (PrP 23-144) (Fig. 3*B*) (58). Y145Stop exhibited a similar phase separation behavior with tau in comparison to full-length PrP, as evident from the turbidity measurements and imaging (Fig. 3*F*, 3*G*, and Fig. S3*C*). Moreover, the turbidity and droplet profile of Nh2-tau:Y145Stop were very similar to that obtained with full-length PrP (Fig. 3*C* and 3*D*). Similarly, as in the case of full-length PrP, with the tau3P variant, Y145Stop showed an increase in turbidity in comparison to the unmodified tau, highlighting the role of electrostatic interactions (Fig. S3*D*). However, the C-terminal globular domain of PrP (PrP 112-231) neither enhanced the phase separation propensity nor the morphological appearance of tau droplets (Fig. 3*F*, 3*G*, and Fig. S3*E*). These findings highlight that the N-terminal intrinsically disordered segment of PrP is the key modulator for complex coacervation of tau and PrP primarily via electrostatic interactions between oppositely charged disordered domains. Previous studies have shown that such domain-specific electrostatic interactions yield nanocomplexes that can serve as primary units for such heterotypic complex coacervates (52). Once created, these units can cohere into a highly dynamic liquid-like dense phase that, although seems homogeneous at the mesoscopic length scale, may contain short-range ordering and dynamic heterogeneity at the nanoscale level. We thus hypothesized that region-specific electrostatic interactions could potentially induce temporal molecular ordering within the condensed phase. Next, we used site-specific picosecond time-resolved fluorescence anisotropy measurements to discern short-range dynamic heterogeneity.

### Electrostatic nanoclusters in heterotypic condensates

In order to investigate the region-specific structural ordering in tau-PrP condensates, we performed fluorescence anisotropy measurements that report the local rotational flexibility. To record the site-specific anisotropy, we used a thiol-active fluorescent dye (Fluorosceine-5-maleimide, F-5-M) to label single-cysteine variants of tau at residue locations 17, 56, 158, 291, 322, 400, and 433 spanning the entire protein chain. Any rise in the steady-state anisotropy is interpreted as the loss of conformational flexibility. Steady-state fluorescence anisotropy values exhibited an increase at each of these locations in tau-PrP heterotypic condensates compared to monomeric dispersed tau as well as tau-only condensates suggesting a dampened rotational motion upon complex coacervation of tau and PrP (Fig. 3*H*). Notably, the anisotropy value at residue 56 located at the negatively charged N-terminal domain of tau showed the most significant increase followed by residue positions at 158 and the 400 (C-terminal end). Additionally, we next measured the changes in the anisotropy of various residue positions in PrP by using respective cysteine variants (Fig. 3*I*). A significant increase in the anisotropy at residue position 99 located at the positively charged intrinsically disordered N-terminal domain of PrP indicated the role of this segment in promoting electrostatically driven complex coacervation of tau and PrP. Moreover, our single-droplet anisotropy measurements for tau and PrP also corroborated our ensemble measurements (Fig. S3*F* and S3*G*). These observations are in line with our results on domain-specific interactions described in the previous section. Together our steady-state fluorescence anisotropy measurements highlighted the central role of the acidic N-terminal segment of tau and the basic N-terminal domain of PrP in forming heterotypic tau:PrP condensates.

Steady-state fluorescence measurements provide time-averaged information and thus cannot distinguish between different modes of rotational dynamics experienced by polypeptide chains. In order to discern the different modes of chain dynamics as well as to estimate the hydrodynamic sizes of the primary units formed via electrostatic interactions, we performed fluorescence depolarization kinetics by picosecond time-resolved fluorescence anisotropy measurements for using fluorescently-labeled tau at the 56^th^ position which showed the most significant increase in the steady-state anisotropy upon complex phase separation of tau and PrP. Monomeric dispersed tau exhibited fast depolarization kinetics that is typical for an expanded polypeptide chain (Fig. 3*J*). A biexponential decay model resolved a fast (sub-nanosecond) rotational correlation time corresponding to the local motion of the fluorophore and a characteristic slower (nanosecond) rotational correlation time that is attributed to collective backbone dihedral rotations and long-range conformational fluctuations. Upon homotypic phase separation of tau, the depolarization kinetics became slower indicating the chain-chain interactions within condensates. Upon heterotypic phase separation of tau and PrP, the depolarization kinetics exhibited an additional slower rotational correlation time (∼ 43 ns) suggesting the formation of heterotypic clusters. Assuming these clusters are spherical and no significant changes in the internal viscosity, the estimated hydrodynamic radius of these tau-PrP heterotypic electrostatic clusters is ∼ 3.6 nm. We would like to point out that this is an approximate estimate of the dimension of the nanoclusters within tau-PrP condensates. Such nanoclusters were detected in other electrostatically driven complex coacervates (32, 52). Taken together, these findings show that domain-specific electrostatic interactions between the acidic N-terminal domain of tau and the basic N-terminal domain of PrP drive the complex phase separation of tau and PrP. Since nucleic acids are known to alter the protein phase behavior, we next set out to examine the effect of RNA on the complex phase separation of tau and PrP.

### RNA drives tau:PrP heterotypic assemblies into multiphasic condensates

The interior of biomolecular condensates is thought to be a dense, yet dynamic, tangled-mesh-like organization of proteins and nucleic acids. RNA, because of its shape, charge, sequence, and conformational plasticity acts as a scaffold for the proteins and introduces multivalency into a multi-component system (59, 60). Because of these properties, RNA can modulate the phase behavior and tune the partitioning and material properties. Typically, RNA-controlled electrostatically driven condensates exhibit a distinct three-regime phase behavior (61, 62). To recapitulate this behavior in the tau:PrP system, we started with individual tau and PrP droplet formation. Tau and PrP demonstrate a reentrant phase behavior, with PrP phase separating over a wider regime than tau for approximately the same protein mass (Fig. S4*A* and S4*B*). This difference in the profile can be attributed to the sequence composition of both proteins; tau is K/G-rich, whereas the N-terminal disordered domain of PrP is enriched with R/G/Y residues. Compared to a K-rich polypeptide, an R-rich polypeptide interacts more strongly with RNA because of its potential to engage with RNA via cation-π interactions in addition to the electrostatic interactions (63, 64). Next, we set out to study the effect of RNA on pre-formed tau:PrP heterotypic condensates. For this ternary system (tau-PrP-RNA) the phase separation regime gets further broadened and taller in comparison to the individual protein-RNA systems (Fig. 4*A*, Fig. S4*A*, and S4*B*). At low RNA concentrations, tau:PrP droplets remain miscible and colocalize within the droplets (type I) (Fig. 4*B*, first panel, and Fig. 4*E*). With increasing concentrations of RNA, however, these assemblies acquire a wide range of immiscible multiphasic morphologies from a core-shell structure to a nested-droplet organization and then a core-shell structure (Fig. 4*B*). This type of architecture is a result of differing interfacial tension, viscosity, and density amongst the interacting coacervates (65-67). In our system, this behavior might be a result of the preferential binding of PrP with RNA through a distinct RNA binding site at the N-terminal domain of PrP, that is absent in tau. In the core-shell morphology (type II) (Fig. 4*B*, second panel, and Fig. 4*E*), at the left side of the RNA-dependent phase diagram, PrP forms the core of the droplet (green) whereas tau distributes itself around the core. With the increasing RNA concentrations, a higher amount of PrP gets recruited inside tau droplets resulting in nested droplets (type III) (Fig. 4*B*, third panel, and Fig. 4E). Interestingly, tau and PrP remain immiscible in this stoichiometry ratio and retain their individual droplet identity as depicted by the FRAP profile for the tau-PrP-RNA ternary mixture (Fig. 4*C*). The formation of this multiphasic nested condensate remained relatively unaffected by changing the order in which the components were added to the reaction mixture suggesting its preference for the tau:PrP:RNA ternary-phase system (Fig. S4*C*). On moving to the right side of the turbidity curve, we again observed a core-shell morphology (type IV), however, with a reverse distribution profile for tau and PrP (Fig. 4*B*, bottom panel, and Fig.4*E*). This observation is further corroborated by our steady-state anisotropy measurements performed for tau:PrP with varying amounts of RNA in the ternary mixture (Fig. S4*D* and S4*E*). This organization can be attributed to the different dissolution concentrations of RNA required for each component because of their net charge difference (63). Inverting the charge on tau:RNA complexes enables them to interact non-specifically with PrP:RNA complexes. The addition of RNA to the pre-formed tau:PrP heterotypic droplets, therefore, results in the switching of coacervate morphology and composition. A further increase in RNA concentration results in the complete dissolution of assemblies. These morphological transitions of these multicomponent condensates appear to be reversible as indicated by the RNA hydrolysis using Ribonuclease A (RNAse A) (Fig. 4*F*, 4*G*, S3, and S4 video). Taken together, our results indicate an RNA-induced tuning of these condensates in a context-dependent manner (Fig. 4*H*). Notably, such interactions can potentially impose an additional level of spatiotemporal regulation in molecular enrichment in condensates. However, the increased enrichment of biomolecules within these condensates makes them vulnerable to aberrant phase transitions into pathological aggregates. Therefore, we next set out to elucidate the effect of tau-PrP complex coacervation on the aggregation propensity of these heterotypic condensates.

**Fig 4.**
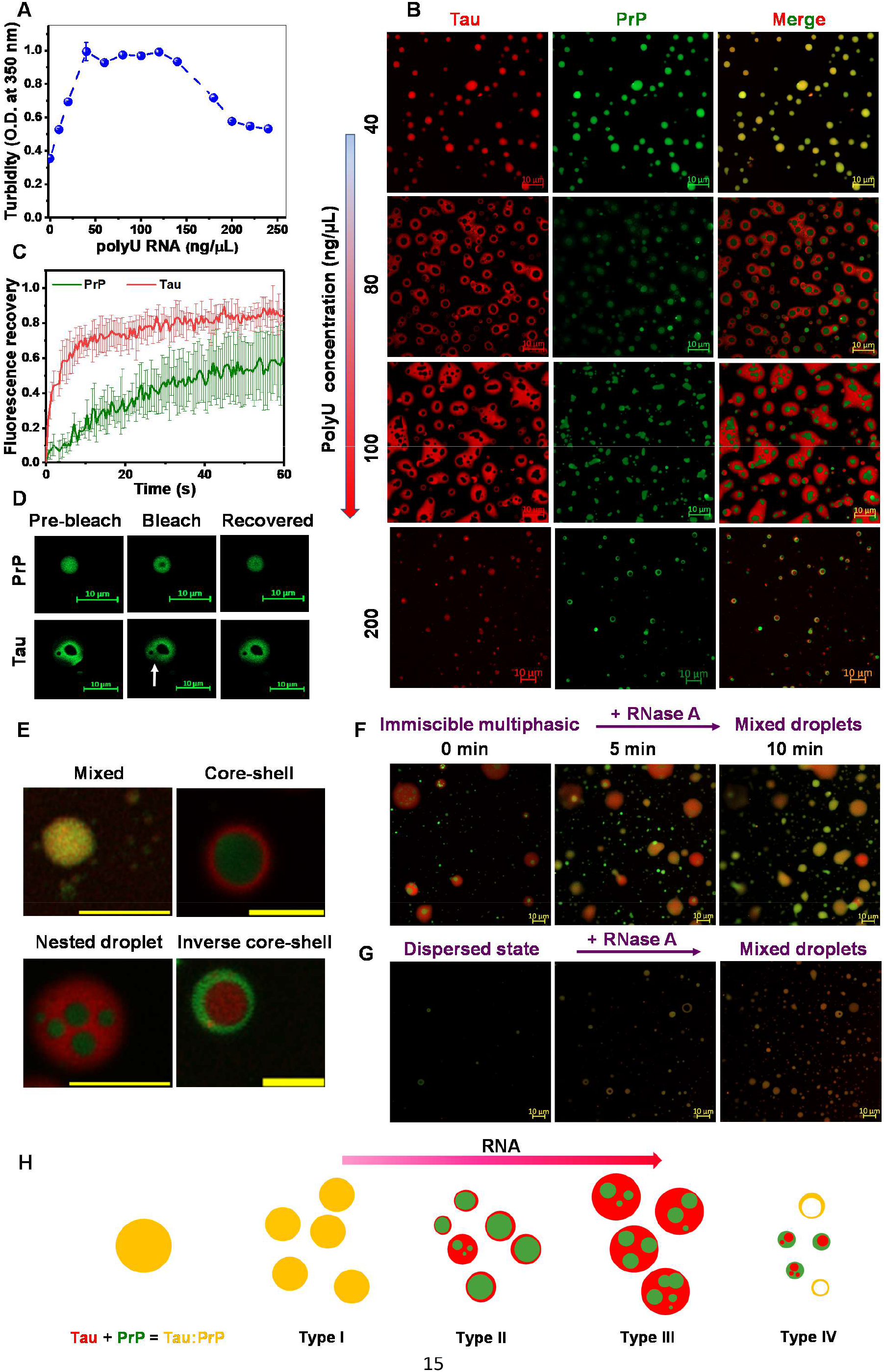
Tau:PrP coacervates form multiphasic condensates in the presence of RNA. **(A)** The Tau:PrP binary system shows a re-entrant phase behavior as a function of RNA concentration. **(B)** Two-color confocal imaging of tau:PrP:RNA (tau-red; PrP-green) ternary system as the function of increasing RNA concentration. With increasing concentration of RNA, tau:PrP coacervates form immiscible multiphasic condensates that transition into a core-shell structure followed by nested droplets. In these multiphasic condensates, PrP concentrates itself in the interior (core) of the tau (periphery) droplet. Further increase of RNA results in inverted core-shell structure (tau forms the core and PrP forms the shell) and then dissolution (scale bar 10 μm). **(C)** FRAP kinetics of tau (red) and PrP (blue) in the tau:PrP:RNA ternary system. Both components retain their individuality in the tau:PrP:RNA mixtures. The data represent mean ± s.d. for n = 3 independent experiments. **(D)** Confocal images during tau:PrP:RNA FRAP experiments. **(E)** Different types of condensates showing mixed, core-shell, nested, and inverted core-shell morphologies obtained in the presence of an increasing concentration of RNA (scale bar of 5 μm is shown in yellow). **(F)** The transition of multiphasic condensates to co-localized mixed droplets in the presence of ribonuclease A (1.5 μM) (scale bar 10 μm). **(G)** Reversible of completely dissolved tau:PrP droplets (in the presence of 400 ng/μL Poly-U RNA) with the addition of ribonuclease A (scale bar 10 μm). **(H)** Simplified schematic of tau:PrP coacervation as a function of RNA concentration.

### Complex coacervates of Tau and PrP convert into solid-like co-aggregates

We observed that upon longer incubation, heterotypic condensates of tau and PrP undergo maturation into gel-like and solid-like aggregates as evident by time-dependent FRAP kinetics over a period of 48 h (Fig. 5*A*, 5*B*). Our FRAP data revealed that tau-PrP droplets undergo a faster transition into a solid-like state compared to tau-only droplets (Fig. 5*A*, 5*B*, and Fig. S5*A*). Next, we asked if these aggregates were amyloid-like. We monitored the conversion kinetics of tau:PrP condensates using a well-known amyloid marker namely, Thioflavin-T (ThT) that exhibits a characteristic emission band at ∼ 483 nm. When compared to tau-only droplets, tau:PrP condensates exhibited a significant increase in the ThT fluorescence after 48 h indicating the formation of ThT-positive aggregates upon the liquid-to-solid phase transition of these heterotypic condensates (Fig. 5*C* and Fig. S5*B*). However, the emission maxima at ∼ 487 nm indicated the presence of amorphous aggregates along with amyloid-like aggregates. We next structurally characterized these phase separation-mediated heterotypic aggregates using vibrational Raman spectroscopy, which allowed us to identify the secondary structural components that were present in these aggregates. For Raman experiments, we used the dense phase of the tau:PrP reaction mixture and monitored the time-dependent changes in the amide I peak (1650 - 1700 cm^-1^) that primarily arise due to the C=O stretching of the backbone. A broad peak spanning from 1660 cm^-1^ to 1675 cm^-1^ indicated the presence of the hydrogen-bonded cross-β amyloid organization as well as amorphous aggregates supporting our ThT-binding results (Fig. 5*D* and 5*E*). Additionally, we used atomic force microscopy (AFM) to visualize these heterotypic aggregates. AFM images of 48-hour-old mixtures revealed the presence of amyloid-like fibrillar morphologies together with amorphous aggregates (Fig. 5*F*). Further, our two-color high-resolution Airyscan imaging of incubated samples indicated the presence of colocalized tau-PrP into rod-like fibrillar structures (Fig. 5*G* and Fig. S5*C*). Taken together, our findings showed that tau and PrP together undergo phase separation into complex coacervates that gradually transition into intermixed aggregates comprising both amorphous and amyloid-like species.

**Fig 5.**
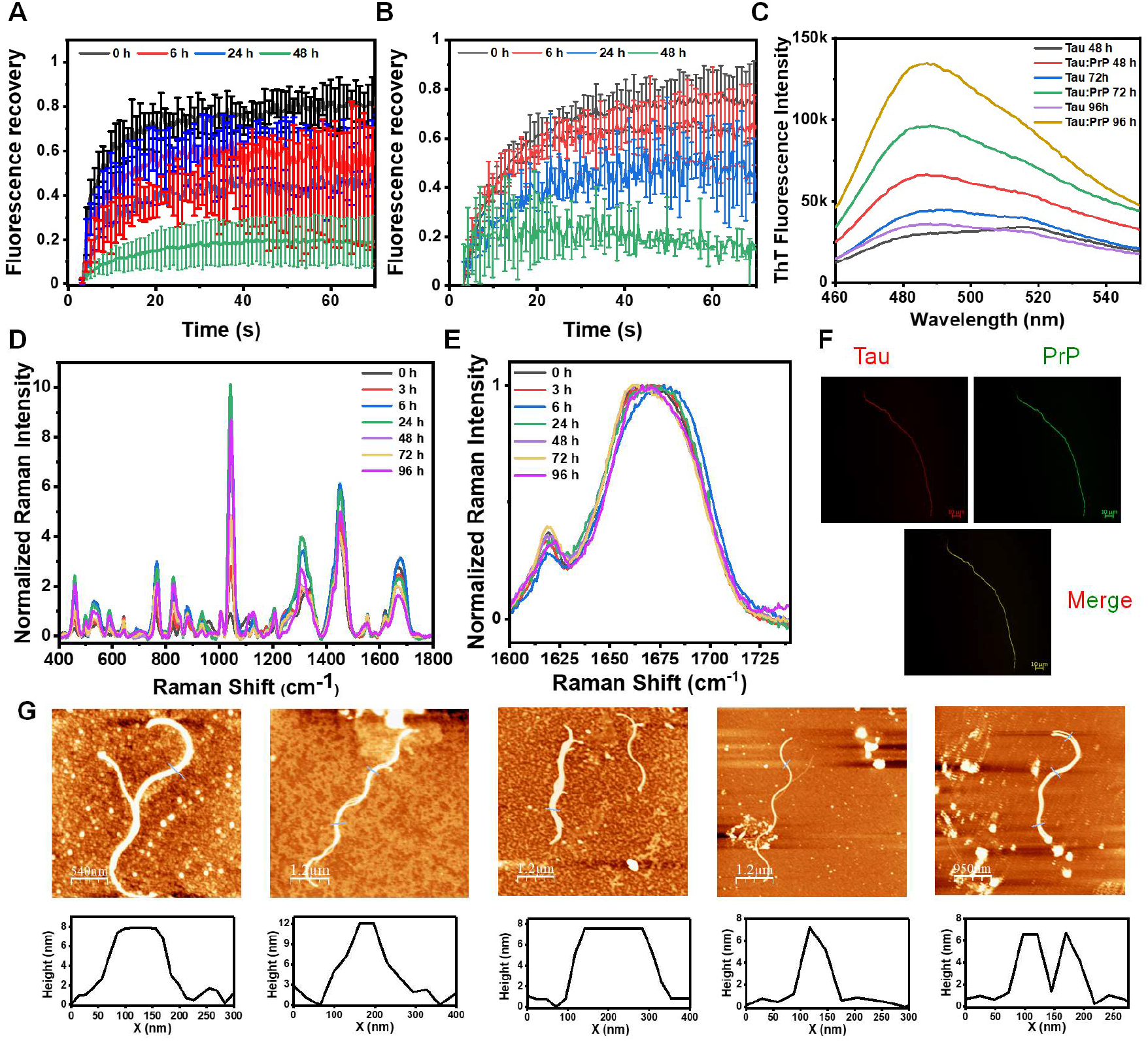
Maturation of Tau:PrP coacervates solid-like aggregates. **(A)** Time-dependent FRAP kinetics of tau and **(B)** PrP inside tau:PrP droplets. Alexa488-labeled protein (∼ 1%) was used in both cases. The data represent mean ± s.d.; n = 3. **(C)** Time-dependent ThT fluorescence spectra of the dense phase of tau-only and tau:PrP reaction mixtures. **(D)** Vibrational Raman spectra of tau:PrP dense phase over time. **(E)** Amide I vibrational Raman band of the dense phase of tau:PrP reaction mixture recorded over time. **(F)** Two-color high-resolution Airyscan image of ∼ 48 h old tau:PrP reaction mixture showing heterotypic association of components, tau (red) and PrP (green), (scale bar, 10 μm). **(G)** AFM images with height profiles of 48 h old tau PrP reaction mixture showing the presence of amyloid fibrils along with amorphous species.

## Discussion

In this work, we showed that tau and PrP undergo spontaneous complex phase separation that is primarily driven by domain-specific electrostatic interactions. Such a complex coacervation gives rise to highly dynamic, two-component, liquid-like droplets. The slower fusion, smaller size, slower internal diffusion, and increased robustness of these droplets formed by the complex phase separation of the two proteins suggested that tau forms condensates comprising a highly networked viscoelastic fluid in the presence of PrP. Moreover, as suggested by our estimated C_sat_ values, the phase separation propensity of tau increases in the presence of PrP (Fig. 1). Tau and PrP interact electrostatically in a domain-specific manner and these heterotypic interactions are further strengthened in the case of the phosphomimetic mutant of tau since glutamic acid residues increase the net negative charge of tau (Fig. 2). By using naturally occurring truncation variants of tau and PrP, we elucidate the importance of their N-terminal domains in driving tau:PrP complex coacervation. Our results provide mechanistic support to such a heterotypic interaction between tau and PrP (68, 69). Our site-specific picosecond time-resolved fluorescence anisotropy data revealed the formation of relatively ordered, short-range, electrostatic nanoclusters of tau and PrP (Fig. 3). Such electrostatic nanoclusters have been shown to act as the primary units of heterotypic condensates (52). These clusters are stable on the nanosecond timescale but can potentially undergo making and breaking on a slower timescale. Such a relay of making-and-breaking of interactions can make the assembly highly dynamic, mobile, and liquid-like as indicated by our FRAP studies. On a slower timescale of FCS and FRAP (milliseconds-to-seconds), this complex coacervate possessing a mobile interior can exhibit a typical liquid-like behavior. In such a liquid-like two-component assembly, a protein can also undergo oligomerization. Our work is in line with the RBP-induced phase separation and vitrification of tau (41). Additionally, our findings also indicate the buffering capacity of RNA (70) in the context of tau:PrP interactions. Tau:PrP heterotypic condensates that remain colocalized and miscible at lower concentrations of RNA assume multiphasic morphologies upon the increase in the RNA concentration. The resulting immiscible multiphasic condensates of differing architecture comprise core-shell and nested droplets reminiscent of nucleolar condensates. These co-existing, immiscible, nested condensates in which the core of the large droplets is constituted by smaller PrP-rich droplets with tau occupying the peripheral regions, arise as a result of differing interfacial tensions between individual condensates formed by the two proteins in the presence of RNA (Fig. 4). This morphology undergoes a transformation into an inverse core-shell and mixed hollow droplets upon the addition of higher amounts of RNA. These morphological transitions of tau:PrP condensates are reversible as evident by RNA hydrolysis by the addition of RNase A. Further, our aging experiments demonstrated that liquid-like tau:PrP condensates gradually mature into solid-like aggregates comprising both amorphous and amyloid-like species. Taken together, our study unveils an intriguing interplay of molecular determinants that promote and regulate the heterotypic phase transition, multiphasic coacervation, and maturation into intermixed ordered aggregates highlighting the molecular basis of overlapping neurodegenerative diseases involving tau and PrP (Fig. 6).

**Fig 6.**
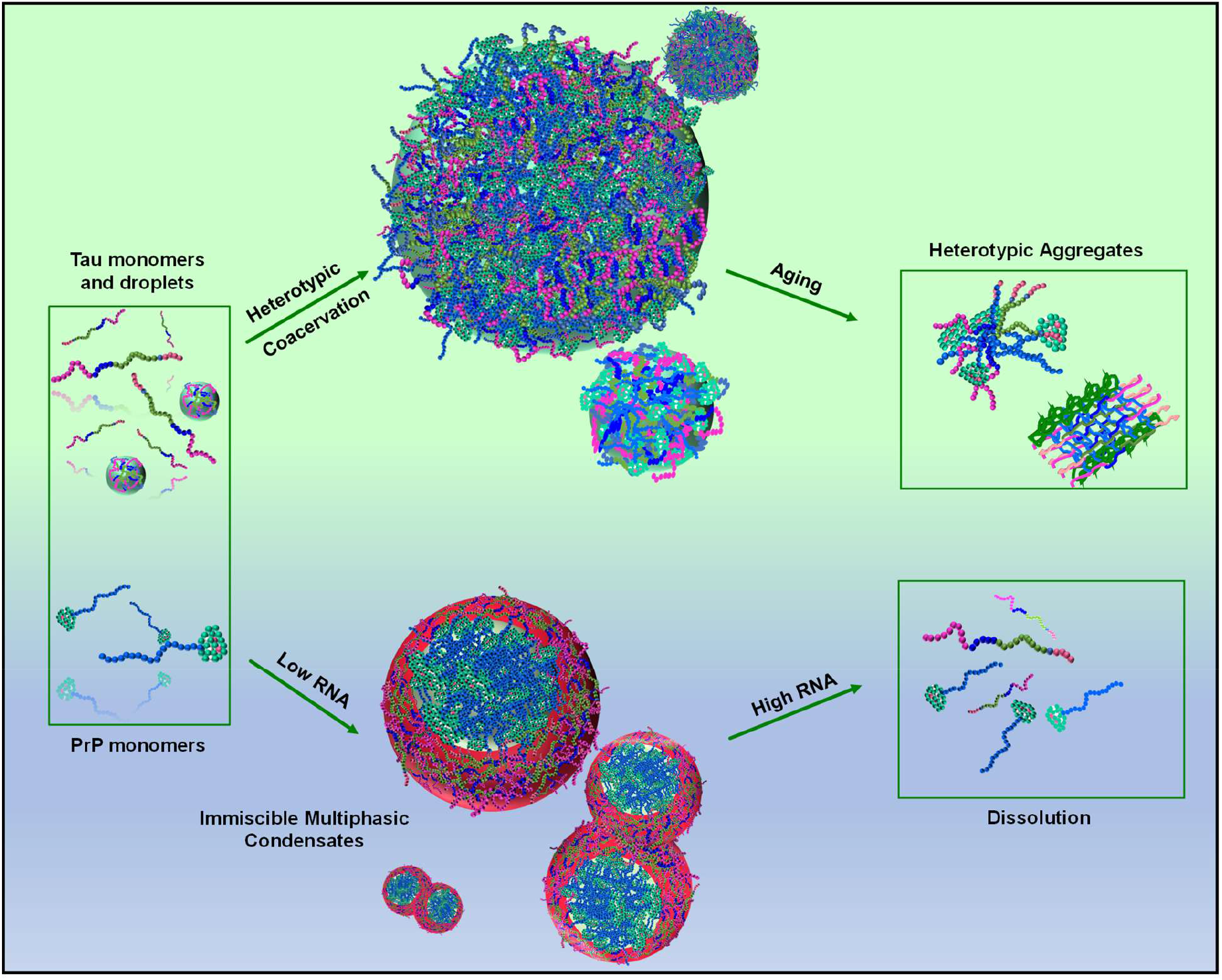
Schematic illustration of complex phase separation of tau and PrP. The N-terminal segments of tau and PrP interact electrostatically to potentiate tau phase separation into heterotypic condensates that mature into intermixed aggregates. Upon the addition of RNA, tau:PrP mixed droplets convert into immiscible, multiphasic condensates and are dissolved at a much higher concentration of RNA.

The inherent sequence attributes of tau, which are also typical of many other phase-separating proteins, regulate its ability to undergo phase separation. Tau, therefore, assembles into membraneless compartments, which sequester tubulin dimer units into well-defined foci, where it is proposed to regulate microtubule polymerization (71). Moreover, tau is also known to localize in other membraneless bodies, including the nucleolus (72) and stress granules (73), where it may contribute to various physiological and pathological roles. Such liquid-like biomolecular condensates of tau can mature into an aggregated form that is a hallmark of neurodegenerative diseases like Alzheimer’s disease and FTLD-associated tauopathies (74, 75). The pathology of tau is not only limited to Alzheimer’s disease and tauopathies but also extends to various forms of prion diseases and synucleinopathies (42-44, 76). The colocalization of tau and PrP as cytoplasmic inclusion bodies and the accumulation of tau NFTs in the brain of prion disease patients (69, 77). Intriguingly, like Aβ oligomers and α-Syn, various forms of tau have been thought to interact with PrP resulting in dysfunction of the synaptic plasticity (76, 78). Although PrP is a GPI-anchored protein, a sparse level of neurotoxic intracellular PrP exists during ER stress. Additionally, in some instances, PrP may only partially cross the ER membrane and adopt one of two transmembrane topologies because of its core hydrophobic region and ineffective translocation (46, 47, 79). In healthy brains of different species, these forms make up no more than 10% of the total PrP molecules, while in transmissible spongiform encephalopathies, they can comprise up to 30%. Two different mechanisms have been proposed for the PrP^C^-mediated neurotoxicity caused by several protein aggregates. In one case, PrP^C^ acts as a receptor and facilitates the internalization of specific extracellular proteins. In another, it activates metabotropic glutamate receptors (mGluRs) by functioning as a transducer to elicit the detrimental effects of certain protein deposits (78, 80, 81). Moreover, PrP^C^-Aβ-oligomer-mediated Fyn kinase activation results in tau hyperphosphorylation (80). Inclusions of hyperphosphorylated tau have also been observed in various acquired and familial forms of prion diseases. Thus, tau and prion show a spectrum of overlapping pathologies following the heterogeneity of these tau-PrP co-deposits. Our results emphasize the phase separation-mediated heterotypic clustering of tau and PrP that can potentially mature into mixed inclusions found in the pathophysiology of several neurodegenerative diseases. Given that variants of tau and PrP are also known to localize in the nucleus (82-85), more precisely in the nucleolar region in the case of tau, our current findings can potentially explain the role of multicomponent tau-PrP assemblies, especially considering the pertinent role of tau, in rRNA-coding, DNA transcription, stabilization, and rRNA processing (86). These types of multicomponent, multiphasic, and anisotropic condensates may be common for many other intracellular nucleoprotein bodies (20, 25). We propose that such interactions might be present in the cellular milieu depending on the subcellular locations. In addition to the direct secretion and absorption of soluble tau isoforms from the membrane-bound receptors, tau, similar to Aβ oligomers and α-syn, is also transmitted amongst neuronal bodies via extracellular vesicles and exosomes (86). These compartments are rich in RNA (87-89) and can therefore provide sites for the formation of tau:PrP:RNA ternary complexes.

In summary, our study relates to both functional and pathological aspects of complex phase separation of tau and PrP with or without nucleic acids. Heterotypic phase separation of α-Syn and PrP has been shown to drive the formation of intermixed α-Syn:PrP amyloids (52). Therefore, PrP can potentially play a central role in recruiting other neuronal IDPs into multicomponent condensates via electrostatic coacervation. Aberrant phase transitions mediated by complex phase separation can potentially serve as a common mechanism for the development and progression of late-life neurodegenerative diseases having overlapping neuropathological features. Targeting such complex phase-separated condensates using small molecules might serve as a potent therapeutic strategy against these debilitating human diseases.

## Acknowledgments

We thank IISER Mohali, Science and Engineering Research Board, Department of Science and Technology (SPR/2020/000333 and CRG/2021/002314 to S.M. and FIST grant # SR/FST/LS-II/2017/97 to the Department of Biological Sciences, IISER Mohali), Ministry of Education, Govt. of India (Centre of Excellence grant to S.M.) for financial support, Prof. Elizabeth Rhoades (University of Pennsylvania, USA) and Prof. Witold Surewicz (Case Western Reserve University, USA) for the kind gift of the DNA plasmids for full-length tau and PrP, respectively, Prof. N. Periasamy (Retd. TIFR Mumbai) for providing us with the fluorescence decay analysis program, Ms. Lisha Arora for helping with the fluorescence decay analysis, and the members of the Mukhopadhyay lab for critically reading this manuscript.

## Authors Contributions

S.K.R. and S.M. conceived the project. S.K.R., R.K., and A.A. performed the experiments and analyses. S.K.R prepared the figures and wrote the first draft. R.K. revised the first draft. S.M. supervised the work, edited the manuscript, obtained funding, and provided the overall direction. All authors discussed the results and commented on the manuscript.

