## Supplementary Information for "Heterotypic electrostatic interactions control complex phase separation of tau and prion into multiphasic condensates and co-aggregates"

#### Materials and Methods

#### Materials

2-(N-morpholino)ethanesulfonic acid (MES), glacial acetic acid, ammonium sulfate, 2-[4-(2-hydroxyethyl)piperazin-1-yl]ethanesulfonic acid (HEPES), sodium phosphate monobasic dihydrate, sodium phosphate dibasic dihydrate, tris base, sodium hydroxide, sodium chloride, imidazole, L-glutathione reduced, 2-mercaptoethanol, Thioflavin T, 1,4-dithiothreitol (DTT), Triton X-100, Thrombin from bovine plasma, PolyU sodium salt, Ribonuclease A (RNase A; from bovine pancreas), phenylmethylsulfonyl fluoride (PMSF), ethylenediaminetetraacetic acid (EDTA), nuclease-free water, were of highest purity grade, obtained from Sigma (St. Louis, MO, USA). Urea and guanidinium hydrochloride were purchased from Amresco. Ampicillin, chloramphenicol, streptomycin sulfate, and isopropyl-β-thiogalactopyranoside (IPTG) were purchased from Gold Biocom (USA). All the fluorescent probes used in this study mainly, fluorescein-5-maleimide (F-5-M), AlexaFluor488-C5-maleimide, AlexaFluor594-C5-maleimide, and IAEDANS (1,5-IAEDANS, 5-((((2-Iodoacetyl)amino)ethyl)amino)Naphthalene-1-Sulfonic Acid), were obtained from Molecular Probes, Invitrogen. SP-sepharose and Ni-NTA resin was purchased from Qiagen. HiLoad™ Superdex-G75 16/600 prep grade (pg), and NAP-10 columns were obtained from GE Healthcare Life Sciences (USA). Amicon membrane filters for concentrating protein were purchased from Merck Millipore. All the buffer solutions were freshly prepared in Milli-Q water and filtered before use. The pH of each buffer solution was adjusted (± 0.02) at 25 °C using a Metrohm 827 lab pH meter.

### Methods

#### Bioinformatic Analysis

Phase separation propensities of tau and PrP were analyzed using FuzDrop/FuzPred (http://protdyn-fuzpred.org/) (1) and catGRANULE (http://s.tartaglialab.com/new_submission/catGRANULES) (2). The distribution of charges throughout both protein chains was analyzed using the Classification of Intrinsically Disordered Ensemble Regions (http://pappulab.wustl.edu/CIDER/analysis) CIDER tool (3). Origin 2020b was used to generate the plots.

#### Site-directed mutagenesis and construct details

All the single cysteine and other variants of full-length human tau, Nh2-tau (26-230), tau truncation (151-399), and tau triple phosphomimetic mutant (tau3P) were created using the tau 6x-Histag-2N4R-17C plasmid which was a kind gift from Prof. Elizabeth Rhoades (University of Pennsylvania, USA). Each variant was made using the QuickChange site-directed mutagenesis kit (Stratagene). The primers used for the respective mutations are listed in Table S1. The 6X-Histidine tag was removed from all these constructs by cloning. The human prion protein (PrP 23-231) plasmid was a kind gift from Prof. Witold Surewicz (Case Western Reserve University, USA). Single cysteine variants of full-length PrP (W31C, W99C, and A120C), its N- and C-terminal truncations, and the cysteine mutants of PrPY145 were created as described before (4). All the mutations were verified by sequencing.

#### Recombinant protein expression and purification

All the variants of full-length tau protein, except Nh2-tau (26-230), were purified in a native condition by using cation-exchange chromatography on an SP-Sepharose column followed by gel filtration on a HiPrep 16/60 Superdex-G-75 (GE) column. Briefly, proteins were expressed by growing bacterial cell cultures at 37 °C, 220 rpm. At O.D.600 = 0.6, expression was induced with 0.5 mM IPTG for 1 h at 37 °C. After harvesting bacterial cells by centrifugation at 4 °C, 4000 rpm for 30 minutes, the cell pellets were dissolved in lysis buffer (20 mM MES, 500 mM NaCl, 1 mM EDTA, 2 mM DTT, 1 mM $MgCl_2$, 1 mM PMSF, pH 6.5). Cells were lysed using a probe sonicator (5% amplitude, 15 seconds on and 10 seconds off pulses, for 25 minutes), following which the lysates were boiled for 10-15 minutes. Cell debris was removed by centrifugation at 11,000 rpm at 4 °C for 30 minutes. The supernatant was collected and treated with streptomycin sulfate (136 μL/mL) and glacial acetic acid (226 μL/mL) to remove any nucleic acid contamination. Again, the residue was removed using high-speed centrifugation. The supernatant was treated with 60% ammonium sulfate, and the precipitated tau protein was collected by high-speed centrifugation for 30 minutes at 4 °C. Protein pellets were dried and dissolved in buffer A (20 mM MES, 50 mM NaCl, 1 mM EDTA, 1mM $MgCl_2$, 2 mM DTT, 1mM PMSF, pH 6.5). The dissolved protein solution was loaded onto the cation exchange column, and the protein was eluted using a linear gradient of 100 % final concentration of buffer B (20 mM MES, 50 mM NaCl, 1 mM EDTA, 1 mM $MgCl_2$, 2 mM DTT, 1 mM PMSF, 1 M NaCl, pH 6.5). Fractions spanning only the peak's central region were pooled together and further polished by gel-filtration in buffer C (25 mM HEPES, 50 mM NaCl, pH 7.4). Purity was ascertained by SDS-PAGE. The purified protein was concentrated using a 10 kDa MWCO Amicon-membrane filter and stored in small aliquots at - 80 °C for future use. For the Nh2-tau, the purification procedure was the same as above, except, in this case, anion-exchange chromatography with a Q-Sepharose column was performed. After precipitation by ammonium sulfate, pellets were dissolved in buffer C. The protein was eluted using a linear gradient of 100% final concentration of salt (25 mM HEPES, 1 M NaCl, pH 7.4). For PrP expression, PrP plasmids were transformed into *E. coli* BL21(DE3) pLysS, and cells were grown in the same conditions described above. Protein expression was induced with 1 mM IPTG at 30 °C for 8 hours. The cells were harvested by centrifugation as described above. The purification for the thrombin cleavable His-tagged constructs of full-length PrP (23-231) and its variants was performed using Ni-NTA chromatography under denaturing conditions along with gradient oxidation (5, 6). Protein was eluted using a buffer containing 500 mM imidazole as described previously. The cysteine mutants of PrP (W31C, W99C, and A120C) were purified under a denaturing condition from inclusion bodies, as mentioned elsewhere. Following purification, the proteins were dialyzed against a phosphate buffer (20 mM sodium phosphate, 50 mM NaCl, pH 6.4). The N-terminal 6xHis-tag was removed by setting up a cleavage reaction using thrombin protease (0.2U/mL) at 37 °C for 5 hours. After the cleavage, the protease was inactivated by 0.2 mM PMSF. Further, to separate the His-tag-cleaved and un-cleaved fractions, the sample was loaded onto a Ni-NTA column and eluted with a 20 mM imidazole buffer (8 M urea, 10 mM Tris-HCl, 100 mM sodium phosphate, pH 8.0). The protein was concentrated using a 3 kDa MWCO amicon membrane filter and refolded (14 mM HEPES, pH 6.8) using a PD10 column. The purity of the protein was confirmed by SDS-PAGE analysis. The concentrations of the proteins were estimated using $\varepsilon_{280}$ = 6400 $M^{-1}cm^{-1}$ for tau full-length, $\varepsilon_{280}$ = 2560 $M^{-1}cm^{-1}$ for both tau truncations, $\varepsilon_{280}$ = 56,590 $M^{-1}cm^{-1}$ for PrP (23- 231), $\varepsilon_{280}$=

43,670 $M^{-1}cm^{-1}$ for Y145 Stop, and $\varepsilon_{280}$ = 14,200 $M^{-1}cm^{-1}$ for PrP (112-231). To avoid freeze-thaw cycles, all the experiments were performed using freshly purified proteins.

**Fluorescence labeling**

Cysteine mutants of tau and PrP were labeled with fluorophores under denaturing conditions at pH 7.4. For fluorescein-5-maleimide (F-5-M) labeling, proteins were mixed in the molar ratio of 10:1 (dye:protein). For Alexa dyes (C5-maleimide), proteins were mixed in a 2:1 molar ratio (dye:protein). 1, 5 -IAEDANS labeling of tau was performed at a 20:1 molar ratio of dye:protein. The reaction mixtures were stirred for 2-3 hours in the dark at room temperature. After completion of the labeling reaction, the excess free dye was removed while buffer exchanging (Buffer C) using a 10 kDa MWCO Amicon membrane filter in the case of tau mutants. In contrast, in the case of PrP mutants, a PD10 column was used. The concentration of the labeled protein was estimated using $\varepsilon_{495}$ = 68,000 $M^{-1}cm^{-1}$, for F-5-M, $\varepsilon_{495}$ = 72,000 $M^{-1}cm^{-1}$, for AlexaFluor488 C5-maleimide, $\varepsilon_{590}$ = 92,000 $M^{-1}cm^{-1}$ for AlexaFluor594 C5-maleimide, and $\varepsilon_{337}$ = 5600 $M^{-1}cm^{-1}$ for 1,5 -IAEDANS.

**Phase separation assays**

Throughout experiments, the concentration of tau protein stock was kept constant at 360 μM. Tau phase separation was induced by diluting the protein (10 μM) in our reaction buffer without salt (droplet buffer:14 mM HEPES, pH 6.8) at room temperature. Tau:PrP droplet formation was achieved by adding tau to PrP in the reaction buffer. The turbidity of the phase-separated samples (tau, tau:PrP, Nh2-tau:PrP, tau:Y145Stop, tau:PrP (112-231), tau:RNA, PrP:RNA, tau:PrP:RNA) was monitored by recording the absorbance at 350 nm, at 25 °C on a Multiskan Go (Thermo scientific) plate reader using 96-well NUNC optical bottom plates. A sample volume of 100 μL was used for these measurements, and raw turbidity data was plotted without background subtraction. The mean and the standard error were obtained from at least three independent sets of measurements performed on a single day. Unless otherwise indicated, tau and PrP concentrations were kept fixed, 10 μM and 20 μM, respectively.

**Confocal microscopy**

Fluorescence imaging experiments were performed at room temperature on a ZEISS LSM 980 Elyra 7 super-resolution microscope equipped with a high-resolution monochrome cooled AxioCamMRm Rev. 3 FireWire(D) camera, using a ×63 oil-immersion objective (numerical aperture 1.4). Less than 1% of respective labeled proteins were doped for imaging droplets with the unlabeled proteins. The freshly prepared samples were incubated for 45 seconds at room temperature. A sample volume of 4-5 μL was placed onto the glass coverslips of a thickness of 1.5 mm. Alexa488-labeled proteins were imaged using a 488 nm laser diode (11.9 mW), and Alexa594-labeled proteins were imaged using a 590 nm excitation source. The images were obtained at a resolution of 1840 × 1840 pixels at 16-bit depth. For the imaging of multiphasic condensates, the required amount of PolyU RNA was added into the pre-formed tau:PrP droplets. In the case of RNase A dependent imaging, 20 μg/mL of RNase was added into the freshly prepared tau:PrP:RNA reaction mixture and 50 μL of the solution was used for imaging. Images were processed and analyzed using the instrument in-built Zen blue 3.2 (3.2) software.

**Fluorescence recovery after photobleaching (FRAP)**

FRAP experiments were performed on the same instrument used for confocal imaging. For all FRAP experiments, Alexa488-labeled (~1%) proteins were used. Measurements were performed for at least three independent samples. For comparison, the best 3-5 traces were used. A region of interest (ROI), having a diameter of 1 μm, was bleached using a 488 nm laser for tau, tau:RNA, PrP:RNA, tau:PrP hetero-protein, and tau:PrP:RNA coacervates. The recovery of the bleached spots was recorded using the in-built ZEN blue 3.2 (ZEISS) software provided with the instrument. Time-dependent FRAP was performed by taking aliquots from reaction mixtures (incubated at room temperature) at desired time points. The fluorescence recovery traces were background corrected, normalized, and plotted using Origin 2020b.

**Estimation of saturation concentration ($C_{sat}$) using sedimentation assays**

The light phase saturation concentration ($C_{sat}$) for reaction mixtures was estimated by ultra-centrifugation. Tau:PrP droplet reactions doped with 10% F-5-M labeled tau were set up and incubated at 25 °C for 5 minutes. The reactions were then ultracentrifuged at ~ 180000 x g at 25 °C for 2 hours. The supernatant was collected and diluted in a salt buffer (14 mM HEPES, 50 mM NaCl) to avoid phase separation. For estimating protein concentration, absorbance at 495 nm was monitored. The resulting value was multiplied by a factor of 10 to obtain the light phase concentration.

**Steady-state fluorescence spectroscopy**

Steady-state fluorescence experiments were performed on a FluoroMax-4 spectrofluorometer (Horiba Jobin Yvon, NJ, USA) using a 1-mm-pathlength quartz cuvette. In all fluorescence studies, the tau and PrP concentrations were kept constant at 10 μM and 20 μM, respectively. For recording the ThT fluorescence, a sample volume of 600 μL was used for all the reactions. Reactions were set up and, at desired time points, pelleted down at 16,400 rpm at 25 °C for 30 minutes. The supernatant was carefully removed, and the obtained pellet was resuspended in a 20 mM sodium phosphate, pH 7.5 buffer. This suspension was further incubated with 20 μM of ThT for 15 minutes before recording the spectrum. The samples were excited at 440 nm, and the emission spectra were collected in the range between 460 nm and 550 nm. For recording the F-5-M-labeled tau and PrP fluorescence (100 nM of labeled protein was mixed with the unlabeled protein), the samples were excited at 485 nm, and the emission spectra were recorded in the range of 510 nm and 600 nm. Steady-state fluorescence anisotropy of F-5-M-labeled tau and PrP were recorded at the emission maximum (~519 nm). The steady-state fluorescence anisotropy ($r_{ss}$) is estimated from the following relationship:

$$r_{ss} = \frac{I_{\parallel} - GI_{\perp}}{I_{\parallel} + 2GI_{\perp}} \quad \text{--- (1)}$$

where $I_{\parallel}$ and $I_{\perp}$ are the parallel and perpendicular fluorescence intensities, respectively, with reference to the excitation polarizer. The G-factor is the geometry factor that was used for correcting the perpendicular components.

**Single-droplet fluorescence anisotropy measurements**

Single-droplet fluorescence anisotropy measurements experiments were performed on a PicoQuant MicroTime MT200 time-resolved fluorescence confocal microscope. A coverslip of 1.5 mm thickness (no. 1) was kept directly on a Super Apochromat 60x water immersion objective with 1.2 NA (Olympus). Laser beams of 488 nm were used for sample excitation and image acquisition. A bandpass emission filter (520/35) for the green dye (F5M and Alexa488) was used before the pinhole. Out-of-focus emission light was blocked by a 50 μm pinhole and the in-focus emission light was then split by a polarizer into 2 detection paths. Single Photon Avalanche Diodes (SPADs) were used as detectors. The correction factor G was estimated using a free dye solution by following manufacturer protocol. Data acquisition and analysis were performed on the commercially available SymphoTime64 software v2.7. For single droplet anisotropy measurements, a region of interest (ROI) selection tool was used for individual droplet's anisotropy value extraction.

**Fluorescence correlation spectroscopy**

FCS measurements were performed on the same instrument as mentioned above (MT200). The confocal volume ($V_{eff}$) and its structure parameter (κ) for our system were determined using a 1 nM solution of Alexa488 which gave us $V_{eff}$ = 1.2 fL and κ = 6.08. These parameters were used as calibration values while curve-fitting data for the monomer and droplets. Dispersed monomeric and droplet solutions (tau and tau:PrP) were prepared by mixing 5 nM Alexa488-labeled tau T17C with unlabeled proteins. The freshly prepared reaction mixtures (50 μL) were spotted onto the coverslip and measurements were performed. In the case of monomer, experiments were performed 50 μm inside the solution, whereas individual droplets were focused in the case of phase-separated solution. Correlation curves (G(t)) were fitted using the triplet model with a 2-diffusion component.

$$G(t) = \left[1 + T\left[e^{\left(\frac{-t}{\tau_{Trip}}\right)} - 1\right]\right] \sum_{i=0}^{n_{Diff}-1} \frac{\rho[i]}{\left[1+\frac{t}{\tau_{Diff}[i]}\right]\left[1+\frac{t}{\tau_{Diff}[i]\kappa^2}\right]^{0.5}} \quad \text{---- (2)}$$

where G(t) is the correlation amplitude, ρ denotes the contribution of the $i^{th}$ diffusing species, T denotes the fraction of the triplet state, $\tau_{Trip}$ is the lifetime of the triplet state, $\tau_{Diff}$ is the diffusion time of the $i^{th}$ diffusing species, and κ is the structure parameter of the corresponding focal volume.

**Time-resolved fluorescence anisotropy measurements**

All ensemble time-resolved florescence anisotropy decays were acquired at 25°C using a time-correlated single-photon counting (TCSPC) setup (Horiba Jobin Yvon, NJ) (7). F5M and IAEDANS labeled proteins were excited using 485 and 375 nm picosecond NanoLED laser diodes. The instrument response function (IRF) was measured using a dilute solution of colloidal silica (Ludox) and the full width at half-maximum (FWHM) was found to be 55 ps. The decay profiles were recorded at the corresponding emission maxima. Using a bandpass of 8nm, the fluorescence intensities were collected at 0° ($I_{\parallel}$) and 90° ($I_{\perp}$) with respect to the geometric orientation of the excitation polarizer. The G-factor, calculated on the basis of free dyes in water, was taken into account to correct the perpendicular fluorescence intensity decays. All measurements were performed for 3 independent replicates, with 3 acquisitions from each sample. The anisotropy decays were analyzed by globally fitting the acquired data according to the following equations:

$$I_{\parallel}(t) = 1/3I(t)[1 + 2r(t)] \quad \text{--- (3)}$$

$$I_{\perp}(t) = 1/3I(t)[1 - r(t)] \quad \text{--- (4)}$$

where *I* denotes the time-dependent fluorescence intensity collected at the magic angle (54.7°). The fast ($\phi_1$) and slow ($\phi_2$) rotational correlation times arising due to the local dynamics of the fluorophore and the segmental dynamics of the backbone, which determine the time-resolved fluorescence decay kinetics can be approximated to a biexponential decay model:

$$r(t) = r_0[\beta_1 e^{\left(\frac{-t}{\phi_1}\right)} + \beta_2 e^{\left(\frac{-t}{\phi_2}\right)}] \quad \text{--- (5)}$$

Here, $r_0$ represents the intrinsic time-zero fundamental anisotropy of the fluorophore. $\beta_1$ and $\beta_2$ denote the amplitudes associated with fast and slow rotational correlation time, respectively. Reduced $\chi^2$ values, the randomness of residuals, and the autocorrelation function gave a measure of the goodness of fit. Anisotropy decays for the dispersed monomers (residue 56 of tau and 99 of PrP) could be fitted using a biexponential decay model. In the case of droplets, a triexponential decay model that also took into an additional slower correlation time ($\phi_3$) was required to describe the time-resolved anisotropy decays as follows:

$$r(t) = r_0[\beta_1 e^{\left(\frac{-t}{\phi_1}\right)} + \beta_2 e^{\left(\frac{-t}{\phi_2}\right)} + \beta_3 e^{\left(\frac{-t}{\phi_3}\right)}] \quad \text{--- (6)}$$

where $\beta_3$ denotes the associated amplitude. A fluorescence probe with a longer lifetime (12 ns) was used to improve the estimation of $\phi_3$, which was estimated to be ~ 43 ns (Table S2). Using these values, the hydrodynamic radii ($R_h$) of the nano-clusters were approximated. The Stokes-Einstein relationship was used for this purpose:

$$\phi_3 = \frac{\eta V}{k_B T} \quad \text{--- (7)}$$

where $\eta$ is the viscosity of the medium, V is the volume of the rotating unit ($V = \frac{4}{3}\pi R_h{}^3$), $k_B$ is the Boltzmann constant, and T is the absolute temperature. The robustness of the recovered correlation time ($\phi_3$) was also assessed by using both free and forced fits.

**Size Exclusion chromatography of tau:PrP complex coacervates**

Multimer formation was also observed using size exclusion chromatography. Here, tau (10 µM) and PrP (20 µM) coacervates were created and incubated at room temperature for suitable amounts of time. Upon completion of incubation, 500 µl of the reaction mixture was run on a

Superedex-G200 10/300 (GE) column, equilibrated with a 20 mM phosphate buffer, pH 6.5. The presence of the protein was observed by measuring absorption at 280 nm. A baseline for the recorded intensities was calculated using the UNICORN software (GE) available with the FPLC setup and subtracted from the observed intensities. The resulting data was exported as a text file and plotted using Origin 2020b. The experiment was repeated thrice with similar observations.

**Raman spectroscopy**

For all Raman measurements, the dense phase from reaction mixtures was used. Reaction mixtures (of 600 µL volume) were pelleted down at specific time points, and the dense phase (pellet) was resuspended in 5 µL of 20 mM sodium phosphate buffer, pH 7.4. The resuspended dense phase was deposited onto a glass slide covered with an aluminum sheet and half-dried (8). An inVia laser Raman microscope (Renishaw, UK) was used for recording all the spectra. The sample was focused using a 100x objective lens (Nikon, Japan), and a 785-nm NIR laser was used for excitation, with an exposure time of 10 s and 100% laser power. Spectra were recorded for tau and tau:PrP droplets at different time points. An edge filter of 785 nm was used for filtering Rayleigh scattering. The Raman scattering was collected and dispersed using a 1200 lines/mm diffraction grating and detected using an air-cooled CCD detector. The instrument's in-built Wire 3.4 software was used for data acquisition. All the data were averaged over 10 scans. Acquired spectra were baseline corrected and smoothened using Wire 3.4. Spectra were plotted using Origin 2018b.

**Atomic force microscopy (AFM) imaging**

AFM images were acquired using an Innova atomic force microscope (Bruker) operating in tapping mode. For sample preparation, 10 µL aliquots were taken from reaction mixtures (incubated for 48 hr at room temperature) and deposited on freshly cleaved, Milli-Q water-washed muscovite mica (Grade V-4 mica from SPI, PA). The samples were incubated for 10-15 minutes at room temperature and were washed with 150 µL of filtered Milli-Q water. The samples were further air-dried using a gentle stream of nitrogen gas before AFM imaging. Data was acquired using NanoDrive (v8.03) software, and the WSxM 5.0D 8.1 software was used for image processing (9). The height profiles were analyzed from WSxM and were plotted using Origin 2020b.

**Table S1.** Primers used for creating point mutations and truncations.

| | |
|---|---|
| 17CT Forward | GGAAGATCACGCTGGGACTTACGGGTTGGGGG |
| 17CT Reverse | CCCCCAACCCGTAAGTCCCAGCGTGATCTTCC |
| S56C Forward | CACTGAGGACGGATGTGAGGAACCGGGC |
| S56C Reverse | GCCCGGTTCCTCACATCCGTCCTCAGTG |
| A158C Forward | CACCGCGGGGAGCATGCCCTCCAGGCCAG |
| A158C Reverse | CTGGCCTGGAGGGCATGCTCCCCGCGGTG |
| S199C_S202E_T205E_ S208E Forward | CAGCTGCCCCGGCGAGCCAGGCGAGCCCGGCGAGCGC TCCCGCACCCC |
| S199C_S202E_T205E_ S208E Reverse | GGGGTGCGGGAGCGCTCGCCGGGCTCGCCTGGCTCGC CGGGGCAGCTG |
| Nh2-tau Forward | ATATATCATATGCAGGGGGGCTACACCATGCACC |
| Nh2-tau Reverse | ATATATCTCGAGTCAACGGACCACTGCCACCTTCTTGG |
| Tau (151-399) Forward | ATATATCATATGATCGCCACACCGCGGGG |
| Tau (151-399) Reverse | ATATATCTCGAGTCACTCCGCCCCGTGGTCTGTCTTGG |
| S291C Forward | GCAACGTCCAGTCCAAGTGCGGCTCAAAGG |
| S291C Reverse | CCTTTGAGCCGCACTTGGACTGGACGTTGC |
| S322C Forward | GAGCAAGGTGACCTCCAAGTGCGGCTCATTAGGC |
| S322C Reverse | GCCTAATGAGCCGCACTTGGAGGTCACCTTGCTC |
| S400C Forward | GTCGCCAGTGGTGTGTGGGGACACGTCTC |
| S400C Reverse | GGAGACGTGTCCCCACACACCACTGGCGAC |
| S433C Forward | GCTGACGAGGTGTGTGCCTCCCTGGCC |
| S433C Reverse | GGCCAGGGAGGCACACACCTCGTCAGC |

**Table S2.** Recovered parameters from time-resolved fluorescence anisotropy decay analyses.

| **Tau-S56C-AEDANS** | **$\phi_1$ ($\beta_1$)** | **$\phi_2$ ($\beta_2$)** | **$\phi_3$ ($\beta_3$)** |
|---|---|---|---|
| Tau monomer | 0.72 ± 0.11 ns (0.59 ± 0.025) | 4.85 ± 0.61 ns (0.41± 0.025) | - |
| Tau droplets | 1.07 ± 0.08 (0.56 ± 0.011) | 8.09 ± 0.11 (0.44 ± 0.011) | - |
| Tau:PrP droplets | 0.45 ± 0.058 (0.39 ± 0.031) | 3.99 ± 0.26 (0.36 ± 0.027) | 43.29 ± 3.63 (0.25 ± 0.021) |

#### Supporting Information Figures

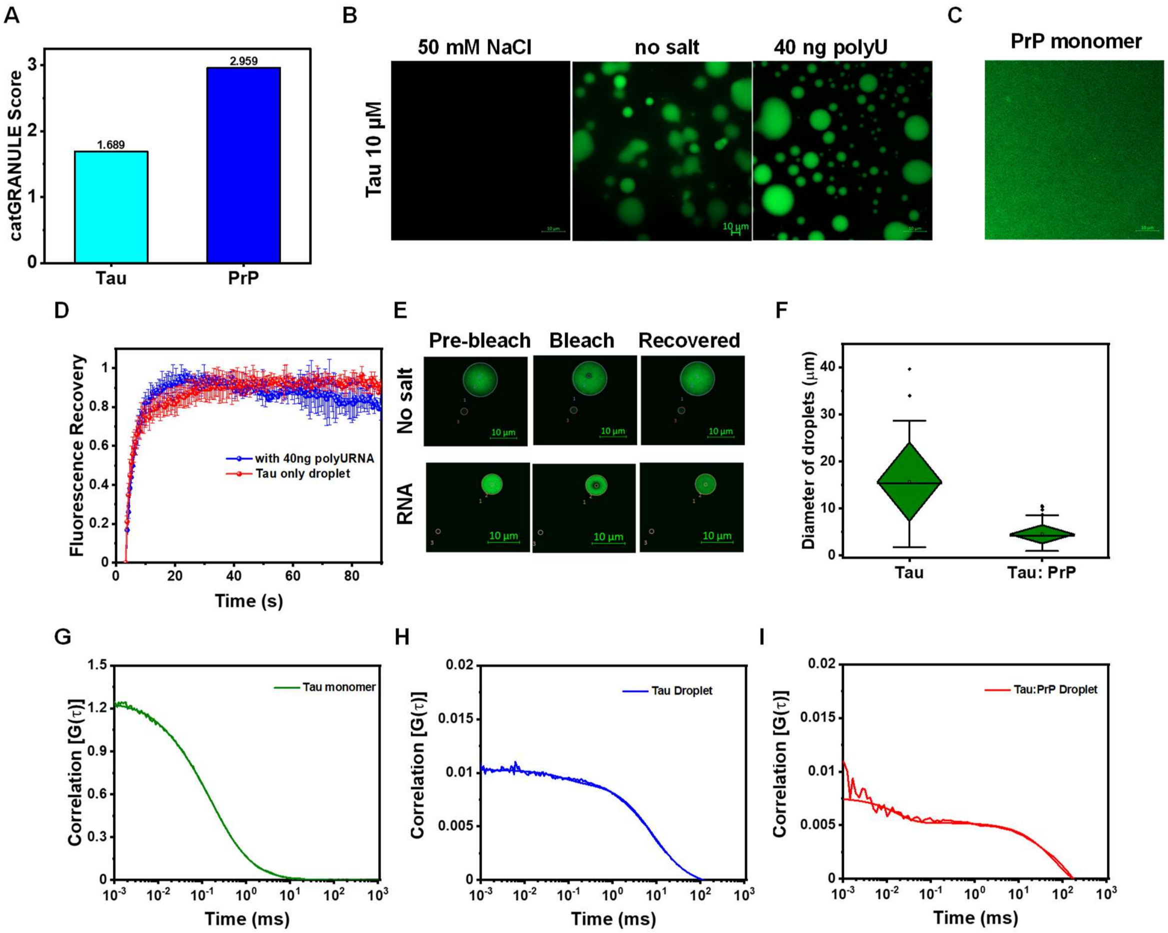


**Fig. S1. (A)** Phase separation propensity of tau and PrP predicted using catGRANULE. **(B)** Confocal imaging of tau monomer (homogeneous phase) in 50 mM NaCl salt buffer (14 mM HEPES), tau-only droplets (10 μM), and tau:RNA droplets. Alexa488-labeled (~ 1%) tau FL17C protein was doped with unlabeled protein for imaging. The imaging was performed at least thrice with similar observations (scale bar 10 μm). **(C)** A mixed homogeneous phase of PrP (monomer). Alexa488-labeled (~ 1%) W99C protein was doped with the unlabeled protein for imaging (scale bar 10 μm). **(D)** FRAP kinetics of tau-only and tau:RNA droplets. Alexa488-labeled protein (~ 1%) was used for FRAP. The data represent mean ± s.d. for n = 3 independent experiments. **(E)** Tau-only (upper) and tau:RNA (lower) droplets images in the time-course of FRAP. **(F)** Comparison of tau-only and tau:PrP droplets size distribution. More than 5 different images were chosen for size analysis. **(G)** FCS autocorrelation plots with fits of tau monomer, **(H)** tau-only, and **(I)** tau:PrP droplets. The normalized versions of these plots are shown in Figure 1I.

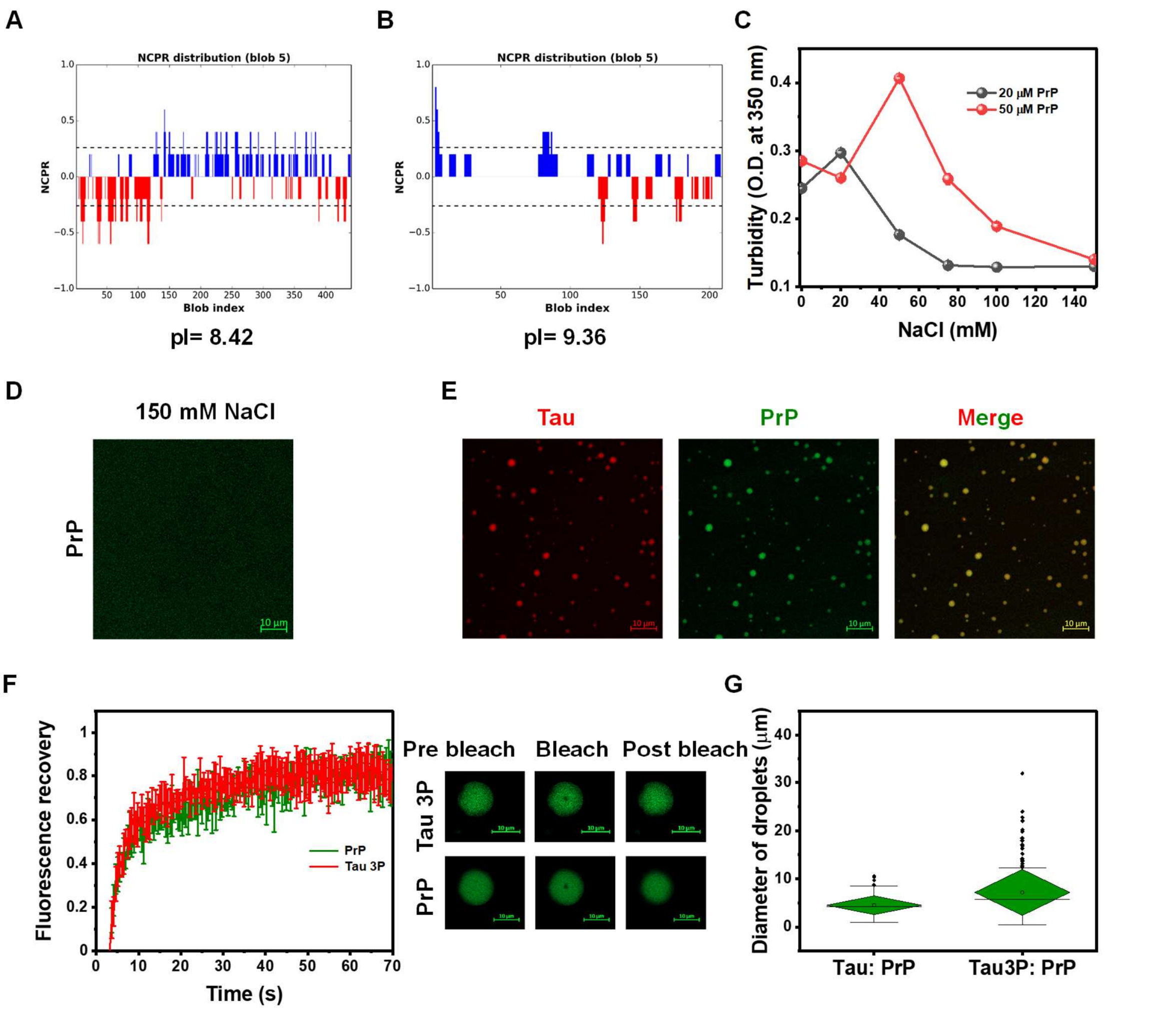


**Fig. S2. (A)** Charge distribution profile (NCPR, net charge prediction) of tau and **(B)** PrP, using CIDER. Red and Blue indicate negatively and positively charged residues, respectively. **(C)** Salt-dependent turbidity (at O.D. 350 nm) of tau:PrP reaction mixtures. The data represent mean ± s.d. for n = 3 independent single-day measurements. **(D)** Confocal images of PrP monomer (50 μM) in the presence of 150 mM NaCl buffer (scale bar, 10 μm). **(E)** At higher concentrations of tau and PrP (30 and 60 μM, respectively) tau:PrP undergo coacervation at physiological salt concentration. **(F)** FRAP kinetics of tau and PrP components in tau 3P:PrP droplets. The data represent mean ± s.d. for n = 3 independent single-day experiments. The adjacent panel shows the images of droplets during FRAP experiments. The imaging was performed at least thrice for all experiments with similar observations. **(G)** Comparison of the size distribution of tau:PrP and tau 3P:PrP droplets. In the case of tau:PrP, the same data set from supplementary figure S1F has been used for comparison.

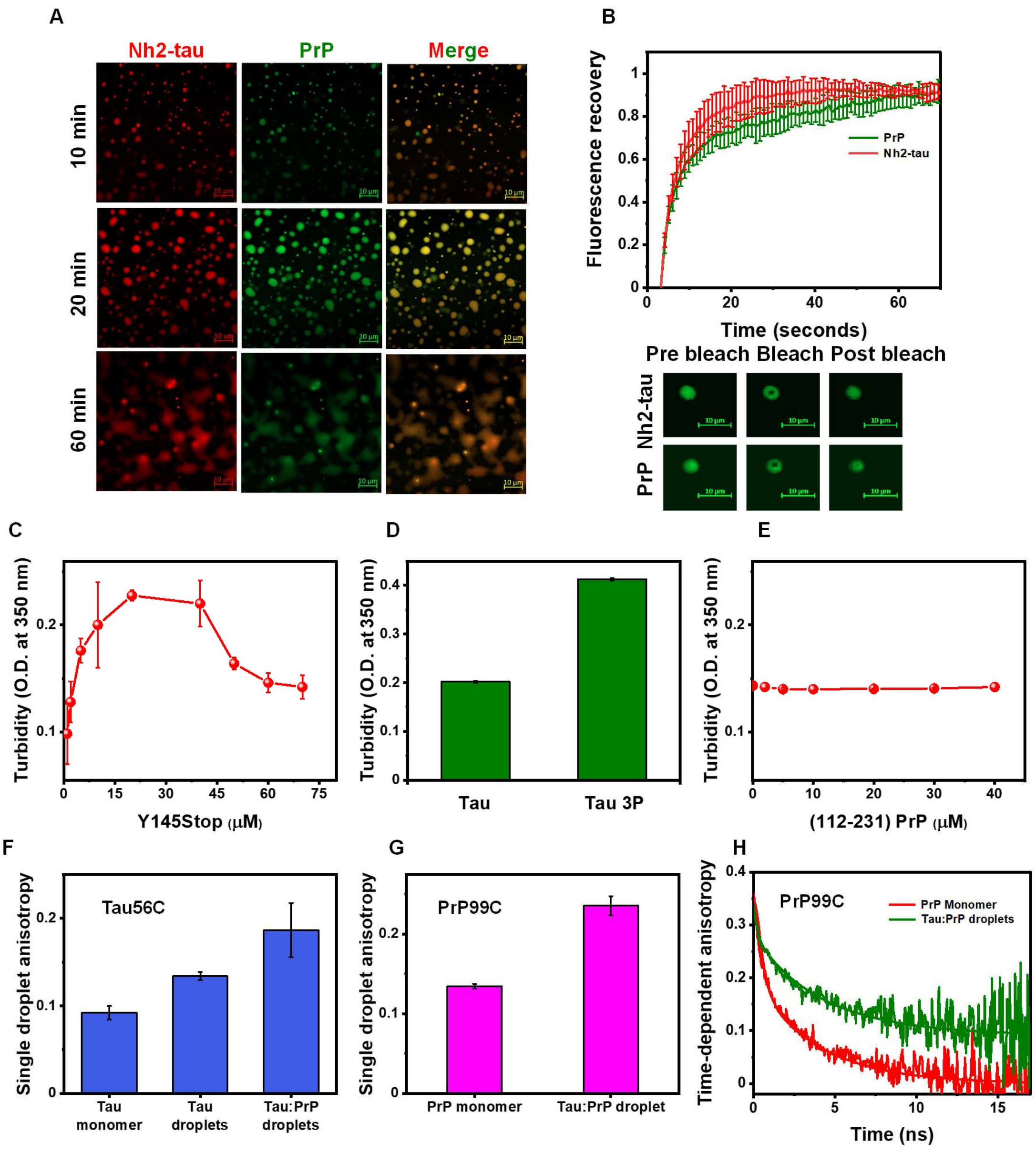


**Fig. S3. (A)** Two-color confocal images of Nh2-tau:PrP droplets after 10 minutes (upper panel), 20 minutes (middle), and 60 minutes (lower panel) (scale bar, 10 μm). These droplets coalesced after 60 minutes. **(B)** FRAP profile of Nh2-tau and PrP components inside, Nh2-tau:PrP droplets. The adjacent lower panel shows the droplet profile during FRAP. The data represent mean ± s.d. for n= 3 independent experiments. **(C)** Turbidity (at O.D. 350 nm) profile of tau:Y145Stop reaction mixtures; with the varying concentration of Y145Stop. **(D)** Comparison of tau and tau 3P turbidity in the presence of Y145Stop. **(E)** No increase in turbidity value was observed for tau:PrP (112-231) reaction mixtures, with the increasing PrP (112-231) concentration. For turbidity measurements, the data represent mean ± s.d. for n = 3 independent single day measurements. **(F)** Single-droplet steady-state fluorescence anisotropy measurements for F-5-M-labeled tau at residue 56, in monomer, tau-only, and tau:PrP droplets. Similar measurements were done for F-5-M-labeled PrP at residue 99 **(G)**. An increase in anisotropy value in phase-separated solution, suggests the domain-specific interaction between tau and PrP. The data represent mean ± s.d. for $n = 3$ independent experiments. Region-specific anisotropy value was extracted from at least $n = 10$ different droplets, in the case of phase-separated solution. **(H)** Time-resolved anisotropy decays of F-5-M labeled PrP W99C in monomer (red) and tau:PrP (olive) droplets. The solid lines are fits obtained using the biexponential and triexponential decay analysis for monomers and droplets, respectively. Refer to Methods, for details of time-resolved anisotropy decay measurements and analysis.

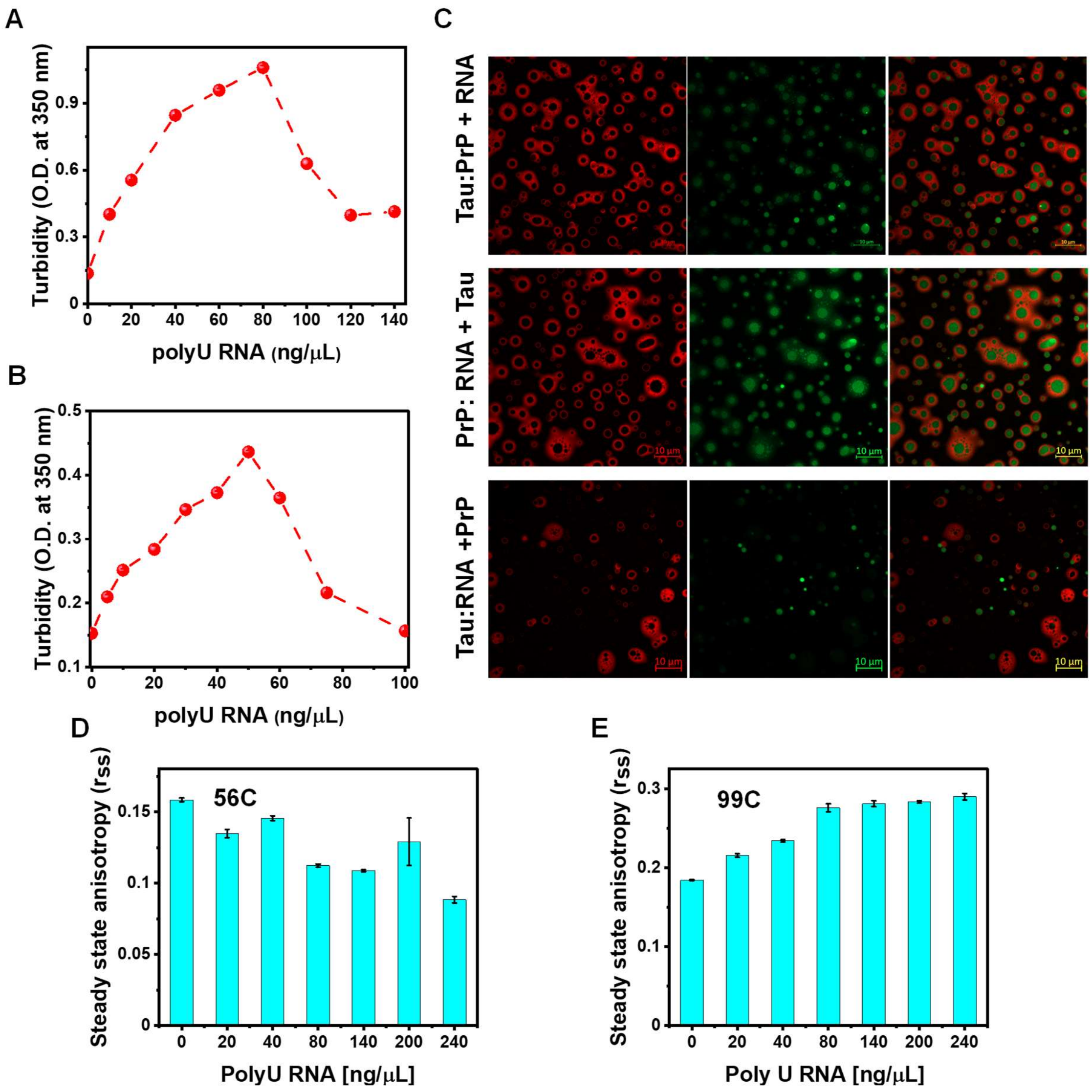


**Fig. S4. (A)** Turbidity (at O.D. 350 nm) profile of PrP and **(B)** tau against increasing concentration of PolyU RNA. **(C)** Two-color imaging of tau:PrP:RNA droplets by changing the order of component addition. Tau was labeled with Alexa594 (red) whereas green labeled (Alexa488) PrP was used (scale bar 10 μm). The imaging was performed at least thrice with similar observations. **(D)** Steady-state fluorescence anisotropy for F-5-M-labeled tau at residue 56 indicates its redistribution within the condensates with increasing RNA concentrations. The data represent mean ± s.d. for $n = 3$ independent experiments. **(E)** Steady-state fluorescence anisotropy for F-5-M-labeled PrP at residue 99 indicates an increase in the order within multiphasic condensates with the increase in the RNA concentration. The data represent mean ± s.d. for $n = 3$ independent experiments.

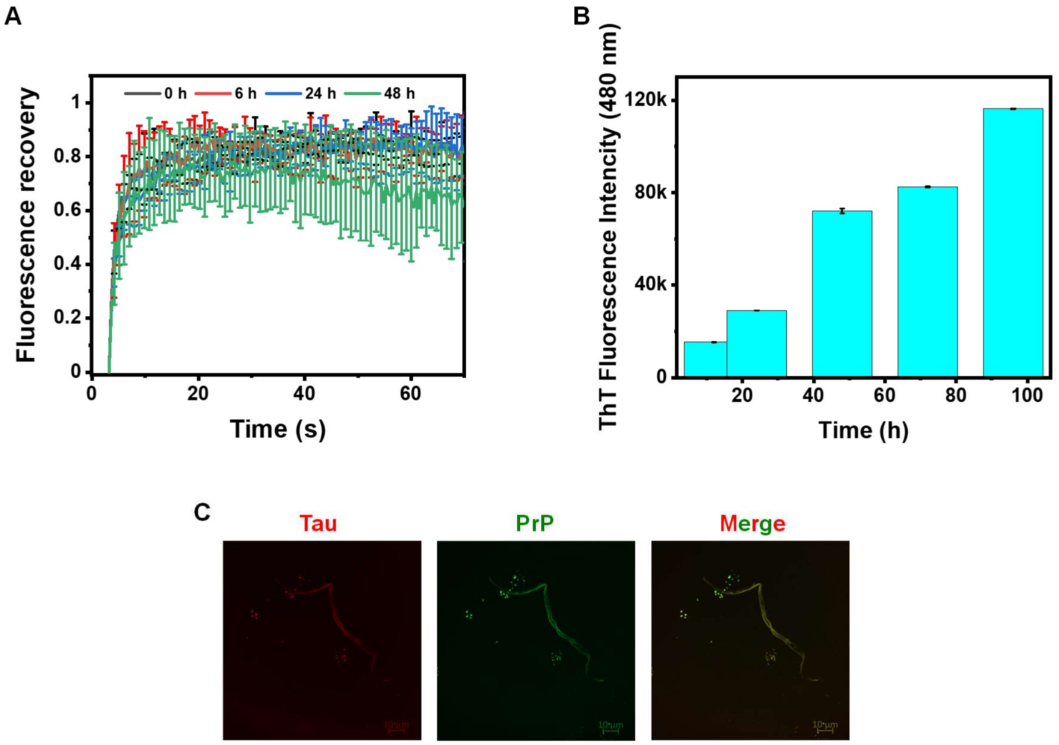


**Fig. S5. (A)** Time-dependent FRAP kinetics of tau-only droplets. Alexa488-labeled (~1% of total protein concentration) labeled (Tau T17C) protein was used for FRAP. The data represent mean ± s.d. for n = 3 independent experiments. **(B)** Single-point measurements of ThT (at 480 nm) fluorescence of tau:PrP dense phase with time. The data represent mean ± s.d. for n = 3 independent experiments. **(C)** Two-color Airyscan image of ~ 48 h old tau:PrP reaction mixture showing the presence of long fibrillar species along with amorphous species in the mixture, tau (red) and PrP (green), (scale bar, 10 µm).

**Movie S1.** Tau-only droplets.

**Movie S2.** Tau:PrP droplets.

**Movie S3.** Multiphasic to mixed tau:PrP droplets in the presence of RNase A.

**Movie S4.** Dispersed phase to mixed tau:PrP droplets in the presence of RNase A.
